# Genetic mutations disrupt the coordinated mode of tyrosinase intra-melanosomal domain

**DOI:** 10.1101/2025.04.21.649833

**Authors:** Sarah Toay, Yuri V. Sergeev

## Abstract

Oculocutaneous albinism type 1 is a genetic disorder caused by the disruption of tyrosinase activity in the melanogenesis pathway. The tyrosinase’s intramelanosomal domain can be subdivided into the catalytic and Cys-rich subdomains, integral for protein stability and catalytic activity. To understand the motions in the tyrosinase intra-melanosomal subdomains and their link to its catalytic activity, we perform essential dynamics on homology models for tyrosinase and the mutant variants R217Q, R402Q, and R217Q/R402Q. Dimensional reduction techniques, such as Principal Component Analysis (PCA), are fundamental to systematically comprehending collective motions in protein structure. The alpha-carbon atomic coordinates for all residues across a 100 ns molecular dynamic trajectory were input into the PCA function, and the results were analyzed alongside correlated movements and free energy profiles for each protein structure. The PCA-identified coordinated movement underlying the stable conformations of wild-type tyrosinase arises within the H9 and H10 helices, which are proximal to the flexible tunnel system and the interface of the catalytic and Cys-rich subdomains. In contrast, genetic mutations R217Q and R217Q/R402Q disrupt the coordinated movement of the tyrosinase intra-melanosomal domain, indicating a cause of mutant variant instability.

**Statement of Importance for Broad Audience:** This study applies molecular dynamics and PCA to the coordinated movements underlying tyrosinase protein stability. The collective motions within the core alpha helix bundle were found in the atomic model of tyrosinase. In OCA1-causing mutant variants, collective motions are lost, suggesting a role of helices in maintaining the stability of wild-type protein. Furthermore, we highlight the proximity of alpha-helices H9 and H10 to the interface of the catalytic and Cys-rich subdomains, which may serve as a potential binding site for targeting by small chaperone molecules to stabilize wild type tyrosinase.

## Introduction

Tyrosinase (TYR) is a principal enzyme in the melanogenesis pathway, producing eumelanin and melanin. Mutations to its gene (*TYR*) are linked to oculocutaneous albinism (OCA), specifically the OCA1 subtype. This genetic disease manifests as hypopigmentation of the skin, hair, and eyes, as well as reduced visual acuity.^1^. The latter may include nystagmus, photophobia, strabismus, and rarely neurodegeneration.^2; 3^. OCA impacts, as a recent review indicates, between 1 in 12,000 to 15,000 people in Europe and 1 in 4,000 and 7,000 in Africa, suggesting higher prevalence than previously recorded^4^. OCA1 is the most prevalent of the subtypes^1; 5^. Phenotypic expression can result from a loss of (OCA1A) or decreased (OCA1B) tyrosinase catalytic activity.^6; 7^.

The tyrosinase protein is constructed of two domains: intra-melanosomal and transmembrane. The former can be further subdivided into the Cys-rich (24-118) and catalytic (179-409) subdomains, which are integral for catalytic activity and protein stability.^8^ The catalytic subdomain contains four alpha-helix bundles (**Figure 1**) at the center of the protein and maintains an active site on one side, while five disulfide bonds stabilize the Cys-rich subdomain.

**Figure 1.**
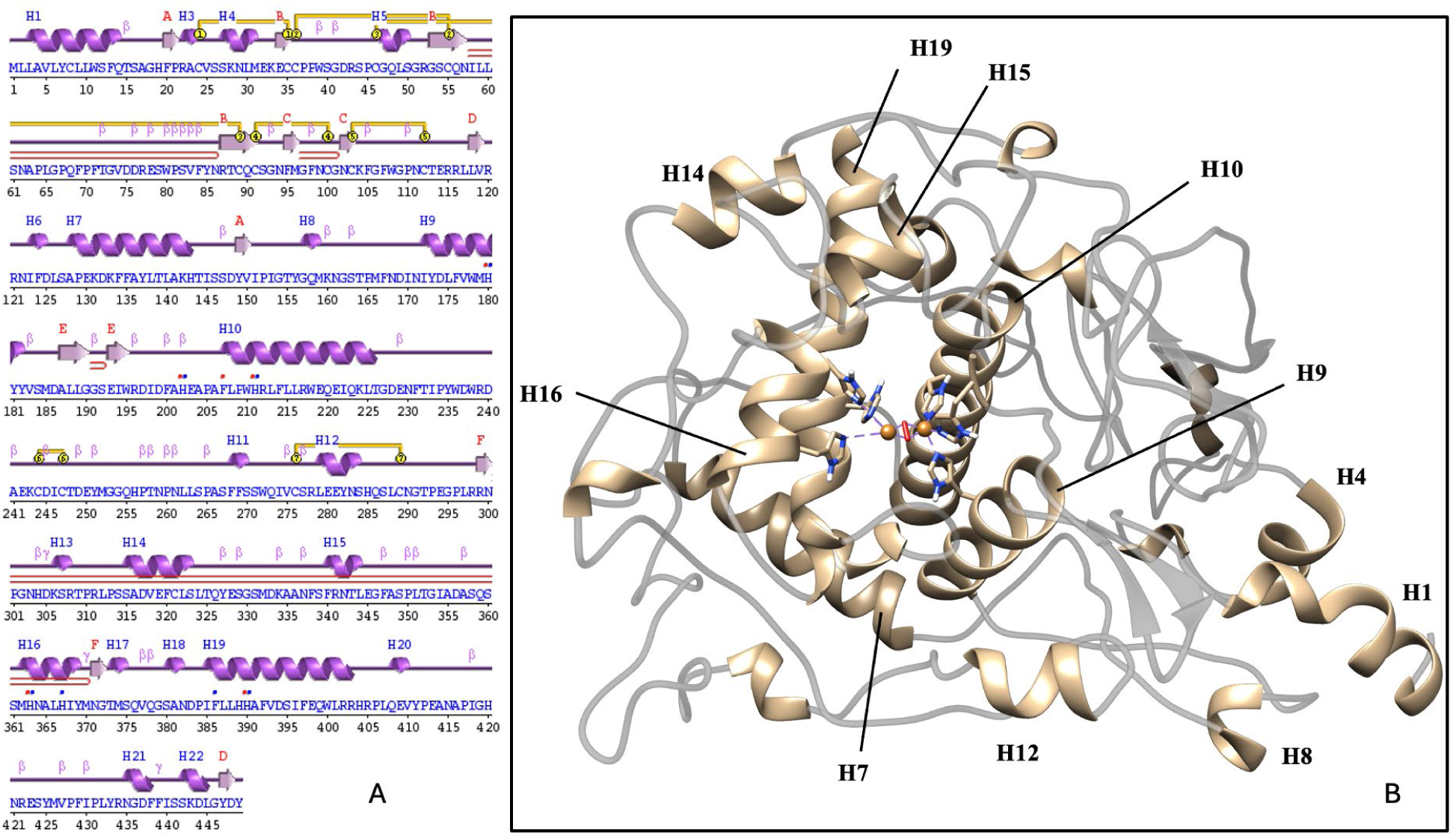
Protein sequence and ribbon structural model of the intra-melanosomal domain of tyrosinase. Amino acid sequence and secondary structure presentation of the tyrosinase intra-melanosomal domain produced using PDBSum generate server (Panel A) and alpha-helical structure (Panel B). Alpha-helices H7, H9, H10, and H19 form a bundle, helping to maintain the tyrosinase catalytic site.

The size of a cavity at the junction of the two subdomains has been shown to correlate with OCA1B mutant stability^9^. The N-terminus consists of a signal peptide, adding 23 residues in length to the tyrosinase protein. Glycosylation has also been shown both experimentally and computationally to provide stability through the N-glycosylation of at least five sites^7; 8; 10^. The active site consists of six histidine residues (180, 202, 211, 363, 367, and 390) coordinating two copper ions and a dioxygen molecule to aid in the oxidation of substrates. No crystal structure of tyrosinase has yet been obtained, so molecular modeling for the protein relies on a Tyrp1-based homology model^8; 11^. Tyrosinase is responsible for completing a series of reactions, converting L-tyrosine to L-3, 4-dihydroxyphenylalanine (LDOPA), and then LDOPA to dopaquinone.^12^.

A variety of tyrosinase activators and inhibitors are known. Ascorbic acid and 2, 3, 5, 4’-tetrahydroxystilbene-2-O-beta-D-glucoside (THSG) are activators, though both can attribute their effects to some degree by acting indirectly through a protein kinase^13-15^. The inhibitors including hydroquinone, kojic acid, and thiamidol, are studied better ^16-18^. One inhibitor, deoxyarbutin, was found to successfully act as a chemical chaperone for a specific tyrosinase mutation, P431L^19^. Chaperones, both molecular and chemical, aid in protein stability, either enhancing the stability of an already folded protein or promoting proper folding^20; 21^. Molecular chaperones have already been recognized as playing roles in tyrosinase folding ^22; 23^. Further, several chemical chaperones have been shown to rescue Tyr activity by promoting proper folding within the endoplasmic reticulum (ER)^24; 25^. The two main types of chemical chaperones are hydrophobic and osmolyte, which reduce improper folding and aggregation by binding exposed hydrophobic regions and minimizing protein movement, respectively^21; 26^. There is also an additional class of chemical chaperones termed pharmacological chaperones, which contain a higher selectivity binding one specific protein^21^. Notable examples of the use of chemical and pharmacological chaperones for treating disease include *N*-octyl-4-epi-β-valienamine (NOEV) for beta-galactosidase deficiency^24^ and large-scale searches to determine potential chaperones for Gaucher and Parkinson’s disease^27; 28^. Small chemical chaperones may be a potential therapeutic route for tyrosinase mutations leading to OCA1.

A previous analysis of the double mutation variant R217Q/R402Q with wild type tyrosinase and corresponding single mutant variants has been completed to describe the basic molecular mechanism underlying the double mutation variant’s phenotypic presentation.^29^. This study combined both computational and experimental techniques to elucidate an important network of interactions between residues R217, F429, and Y149 disrupted in the R217Q and R217Q/R402Q variants. This combined methodology is useful for a targeted analysis but cannot comprehensively and systematically describe the macro-level dynamics within tyrosinase and mutant variants. Rather, dimensional reduction techniques such as principal component analysis (PCA) and normal mode analysis (NMA) are appropriate to describe collective motions^30^. PCA has not been used to study the protein dynamics of tyrosinase or the structural differences between wild-type and mutant variants. Thus, we chose to apply PCA in our dynamic analysis of wild type tyrosinase and the mutant variants R217Q, R402Q, and R217Q/R402Q, of which the molecular mechanism of pathogenicity has previously been explored. The simultaneous comparison of this tetrad of structures provides insights into the large-scale, coordinated movements of tyrosinase, the polymorphic R402Q mutant variant, and the complexities of multiple mutation variants alongside its constituents.

Principal component analysis (PCA) is a useful dimension-reduction technique for handling large datasets^31^. Its use in protein dynamics is well documented, particularly for determining ‘essential dynamics’, or large-scale movements^32-34^. Essential dynamics is often performed on the x, y, and z coordinates of the alpha carbons of each residue within the system ^32; 35; 36^. To investigate additional protein states, trajectory concatenation can be applied to provide information into a larger array of subspaces^37^. Recently, it has also been used extensively in conjunction with molecular docking to determine potential inhibitors^38-40^. It can be used together with dynamic cross-correlation matrices (DDCM) and porcupine plots. DCCM plots display coordinated movements between different residues in a system, while porcupine plots visualize the residues’ movements with an arrow, providing greater insights into the comprehensive dynamic movements of protein systems^38; 39; 41; 42^.

Free energy landscapes for tyrosinase or mutant variants are not available. Previous tyrosinase mutant studies have provided Gibbs free energy changes ΔΔG using FoldX^9; 43^ which one study has compared to experimentally determined values using temperature-dependent kinetics^44^. While informative in the basic understanding of mutation effects on tyrosinase structure and decisive in its singular numeric form, it lacks nuance, only providing a snapshot of one energy state rather than the often-complex array of local and absolute minima and maxima that may be viewed with a free energy landscape. The latter allows for the visualization of protein energy states, both stable and transition, providing a more accurate representation of protein systems, making free energy landscapes a common practice^45-47^. Additionally, no computational energy analysis of either method has been completed for a multiple mutation tyrosinase variant. Without this, a comprehensive understanding of the structural and catalytic effects of the interaction of multiple mutations in one protein structure cannot be achieved.

Therefore, in this study, we perform essential dynamics on homology models for tyrosinase and the mutant variants R217Q, R402Q, and R217Q/R402Q to describe the movements and stability of wild type tyrosinase and how each mutation or set of mutations influences the protein stability. Through PCA, DCCM, porcupine, and tunnel analyses, we show flexibility at the interface of the Cys-rich and catalytic subdomains for wild type tyrosinase. This flexibility makes the interface a natural site for mutations to generate instability in disease-causing mutations, such as the R217Q and R217Q/R402Q mutant variants in which subdomain separation occurs. The interface, lined with residues of N- and C-termini, may correlate with an additional Western Blot band for each of the mutant variants, missing approximately 127 residues. The flexibility also lends this junction as a potential location for therapeutic stabilization with chaperone molecules, to which we supply several binding site suggestions. Finally, we report the first free energy landscape for tyrosinase and selected mutant variants, providing evidence for a complex set of both stable and unstable energy states. This landscape, alongside computationally calculated ΔΔG values, is also described for the first time for a multiple mutation variant, R217Q/R402Q.

## Results

### Molecular Dynamics

To understand the basic protein behavior of the molecular dynamic trajectories, three properties were monitored throughout the trajectory – root mean square fluctuation (RMSF), solvent-accessible surface area, and radius of gyration. RMSF provides insights into the movement of each residue over time. A higher magnitude of RMSF signifies that a residue is displaying more movement and thus might increase instability. The average RMSF for the alpha carbon atom of each residue was projected onto tyrosinase and mutant variant PDB structures to show differences in the areas undergoing movement (**Figure 2A-D**).

**Figure 2.**
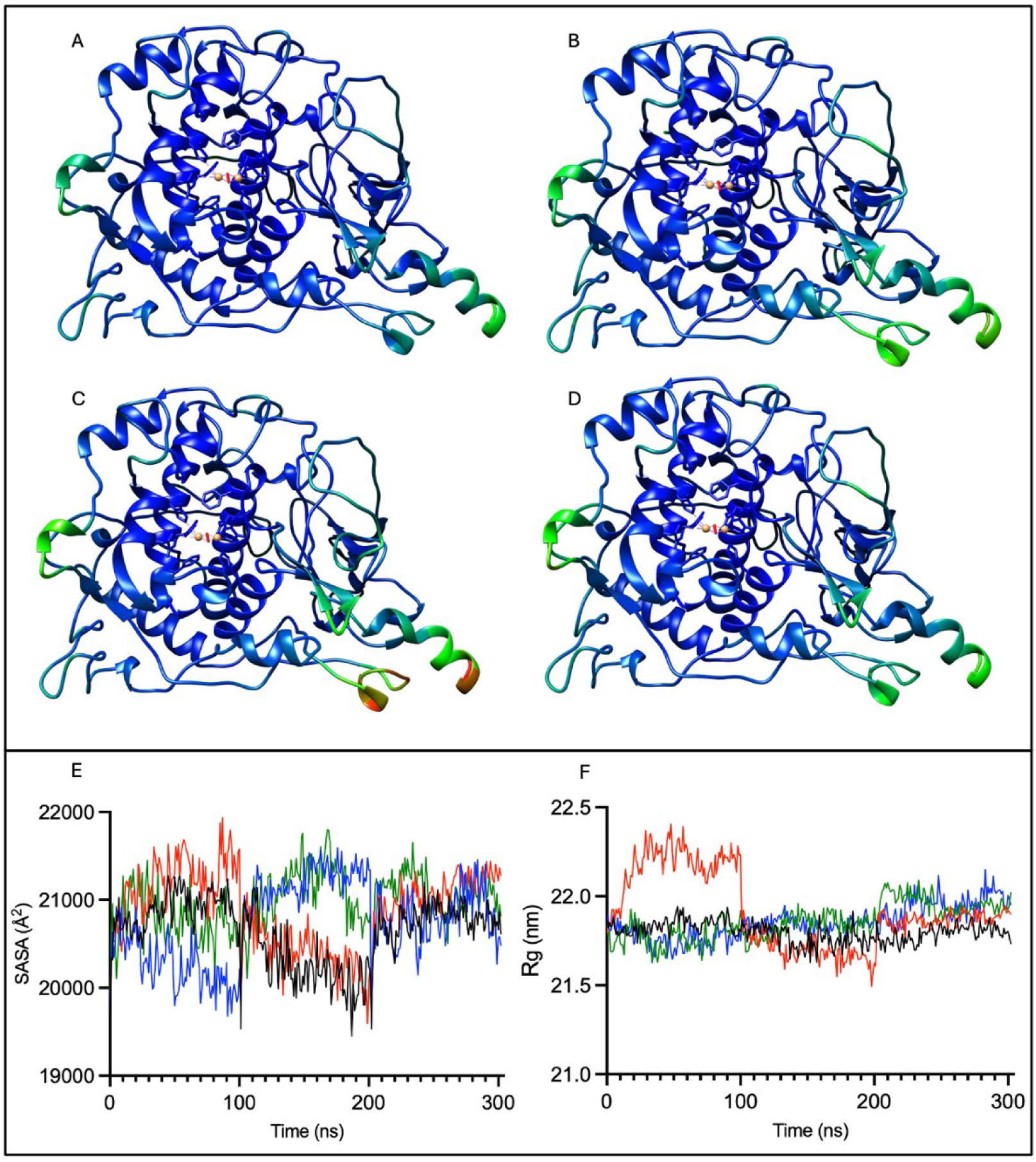
RMSF colored tyrosinase structures and changes over time for solvent-accessible surface area and radius of gyration. The RMSF for tyrosinase (Panel A), R217Q (Panel B), R402Q (Panel C), and R217Q/R402Q (Panel D) are colored with a rainbow scale with larger values in red and smaller values in blue. The concatenated trajectories for SASA (Panel E) and Rg (Panel F) are colored by protein: wild type Tyr (black), R217Q (red), R402Q (green), and R217Q/R402Q (blue) proteins.

All four protein structures were colored on the same rainbow scale with a minimum RMSF value of 0 Å, depicted in red, a middle value of 2.12 Å, shown in bright green, and a maximum RMSF value of 4.23Å, represented in dark blue. Relatively little movement occurs within most of the protein, as values fall near the RMSF minimum. The major movements exist within the signal peptide region, the previously analyzed residues V150-I170 (H8 helix) region^29^, the hairpin turn in the beta sheets, and the N300-L312 region. The R217Q mutant variant also displays increased movement in the residues N57-F71 and C35-G41, colored in lighter shades of blue and green. There are more nuanced differences between wild-type tyrosinase and the double mutant variant in regions R403-A414, W236-D249, C35-G41, and N57-F71. The R402Q also displays a slightly elevated movement in the C35-G41 and N57-F71 regions. Therefore, in addition to previously described movements^29^, 3D-visualization of RMSF highlights three additional regions prone to movement.

Additionally, the solvent-accessible surface area and radius of gyration were calculated for each concatenated trajectory (**Figure 2E-F**). The former provides insights into the total amount of the protein’s surface that can interact with solvent. A higher SASA may suggest a protein’s increased propensity for unfolding as opposed to proteins with a lower SASA where internal, hydrophobic residues remain embedded within the protein’s core. Concurrently with SASA, Rg was calculated to provide insights into the compactness of protein structure. Rg describes the spread of the protein’s atoms about its center of mass. A larger Rg magnitude indicates that a protein is less compact. Descriptive statistics for both SASA and Rg are available (**Supplementary Table S1 and Table S2**).

All four proteins exhibit a distribution of SASA values within approximately a 2000 Å^2^ range (Figure 2 E-F). The location within this range distinctly varies by trajectory, which differs solely in random seed. Trajectories 1 and 3 of the wild-type protein exhibit a slightly higher SASA and Rg than trajectory 2. The R217Q mutant variant follows a similar pattern. While most of the Rg value distribution for all four proteins is within approximately a 0.5 nm range, the first trajectory for the R217Q mutant variant displays a notably higher range. This trajectory is the cause for the 0.5 nm extended range of this mutant variant. It is important to note that smaller differences in Rg, though present, may visually appear minimal due to scaling. The R402Q and R217Q/R402Q mutant variants display low SASA and Rg in the first trajectory and then higher magnitudes of both in the second and third trajectories. There are no significant differences between mean SASA and Rg values for different protein variants. Therefore, we see subtle contrasts between various trajectories for a single protein and among concatenated patterns between the four proteins, demonstrating variability across time and between structures.

### Principal Component Analysis

Principal component analysis (PCA) was completed for each of the protein structures of tyrosinase and mutant variants (**Table 1**).

**Table 1.** Principal Components Summary.

| Structure | PC1<br>Eigenvalue | PC1 Cumulative Proportion<br>of Variance (%) | PC2<br>Eigenvalue | PC2 Cumulative Proportion<br>of Variance (%) | PC3<br>Eigenvalue | PC3 Cumulative Proportion<br>of Variance (%) |
| --- | --- | --- | --- | --- | --- | --- |
| WT | 252.3 | 18.73 | 148.0 | 29.71 | 64.04 | 34.47 |
| R217Q | 287.0 | 21.31 | 224.0 | 37.94 | 69.33 | 43.09 |
| R402Q | 247.0 | 18.34 | 161.7 | 30.34 | 97.70 | 37.59 |
| R217Q/R402Q | 296.1 | 21.98 | 168.4 | 34.48 | 87.32 | 40.96 |

The principal components (PCs) to be analyzed were determined using a scree plot (**Supplementary Figure S1**) alongside the summary data. PCs with the largest eigenvalues were chosen before leveling off and an attempt was made to maximize cumulative variance. The cumulative variance in the first 3 PCs was 35-45%. As compared to recommended and recent field applications, a soft threshold of 70%^31^ and a range from 45-60%, respectively^38-40^. This may be accounted for by the large nature of the protein system, standing at 449 residues in total. Given the magnitude of the system and biochemically relevant results, we determined the cumulative variance to be sufficient.

Next, each score plot for PC1, PC2, and PC3 was analyzed using silhouette analysis and K-means clustering to systematically determine the number of clusters and time points were associated with each (**Figure 3, Supplementary Table S3)**. All four proteins exhibit three clusters. The PC3 dimension failed to produce substantial, meaningful results as variation tended to lie in the planes of PC1 and PC2.

**Figure 3.**
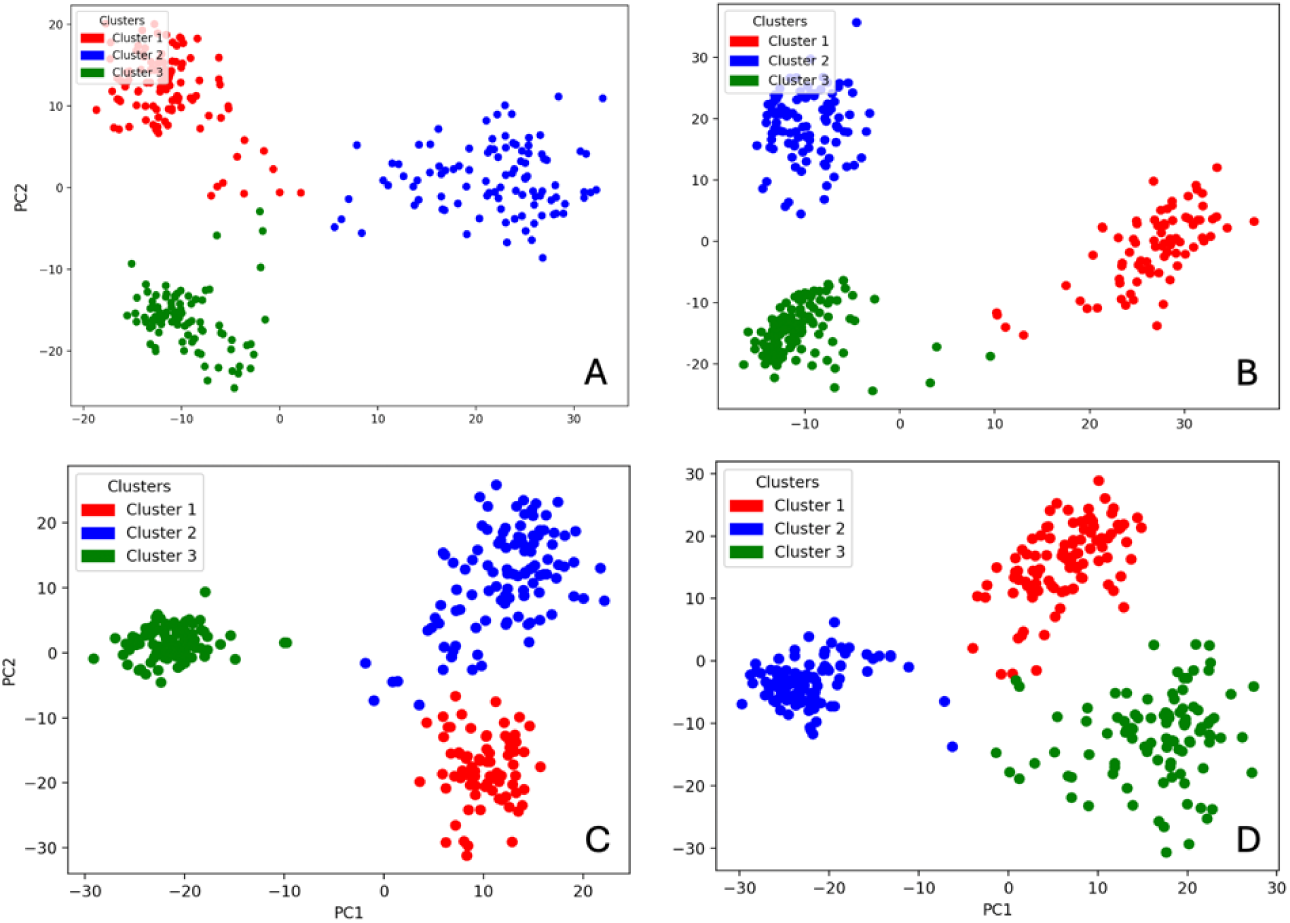
K-means score plots for principal components. Presented are PC1 and PC2 of WT (Panel A), R217Q (Panel B), R402Q (Panel C), and R217Q/R402Q (Panel D).

For all four proteins, selected clusters are roughly associated with a trajectory exploring different conformational subspaces, with mild overlapping between trajectories. Each cluster can be associated with a specific range of timestamps: WT Cluster 1 (1-103, 204-215 ns), WT Cluster 2 (104, 106-203 ns), WT Cluster 3 (105, 216-303 ns), R217Q Cluster 1 (16, 19-101 ns), R217Q Cluster 2 (106-202 ns), R217Q Cluster 3 ( 1-15, 17-18, 102-105, 203-303 ns), R402Q Cluster 1 (22-100 ns), R402Q Cluster 2 ( 1-21, 102-201 ns), and R402Q Cluster 3 (0, 101, 202-303 ns), R217Q/R402Q Cluster 1 (1, 3-100, 202 ns), R217Q/R402Q Cluster 2 (2, 102-201, 203 ns), R217Q/R402Q 3 (0, 101, 204-303 ns). Timestamps 1, 102, and 203 are the 0 ns timeframes for each trajectory, respectively.

In Tyr, Clusters 1 and 3 share characteristics as defined by negative PC1 subspace, but differ by positive and negative PC2 subspaces, respectively. Cluster 2 can be characterized according to the positive PC1 subspace. In the R217Q mutant variant, Clusters 2 and 3 share characteristics as defined by negative PC1 subspace, but differ by positive and negative PC2 subspaces, respectively. Cluster 1 can be characterized according to the positive PC1 subspace. In the R402Q mutant variant, Clusters 1 and 2 share characteristics as defined by positive PC1 subspace, but differ by negative and positive PC2 subspaces, respectively. Cluster 3 can be characterized according to the negative PC1 subspace. In the R217Q/R402Q mutant variant, Clusters 1 and 3 share characteristics as defined by positive PC1 subspace, but differ by positive and negative PC2 subspaces, respectively. Cluster 2 can be characterized according to the negative PC1 subspace.

Next, the top PC loadings were analyzed in two methods. First, the top overall loadings as defined by a threshold of 0.7 were reflected onto reference PDB structures to highlight the structural correlates to each PC subspace (**Supplementary Figure S2, Supplementary Paragraph S1**). Then the top ten percent of loadings aligning with each cluster were extracted, utilizing the Euclidian distance from the origin and K-means clustering (**Figure 4, Supplementary Figure S3**).

**Figure 4.**
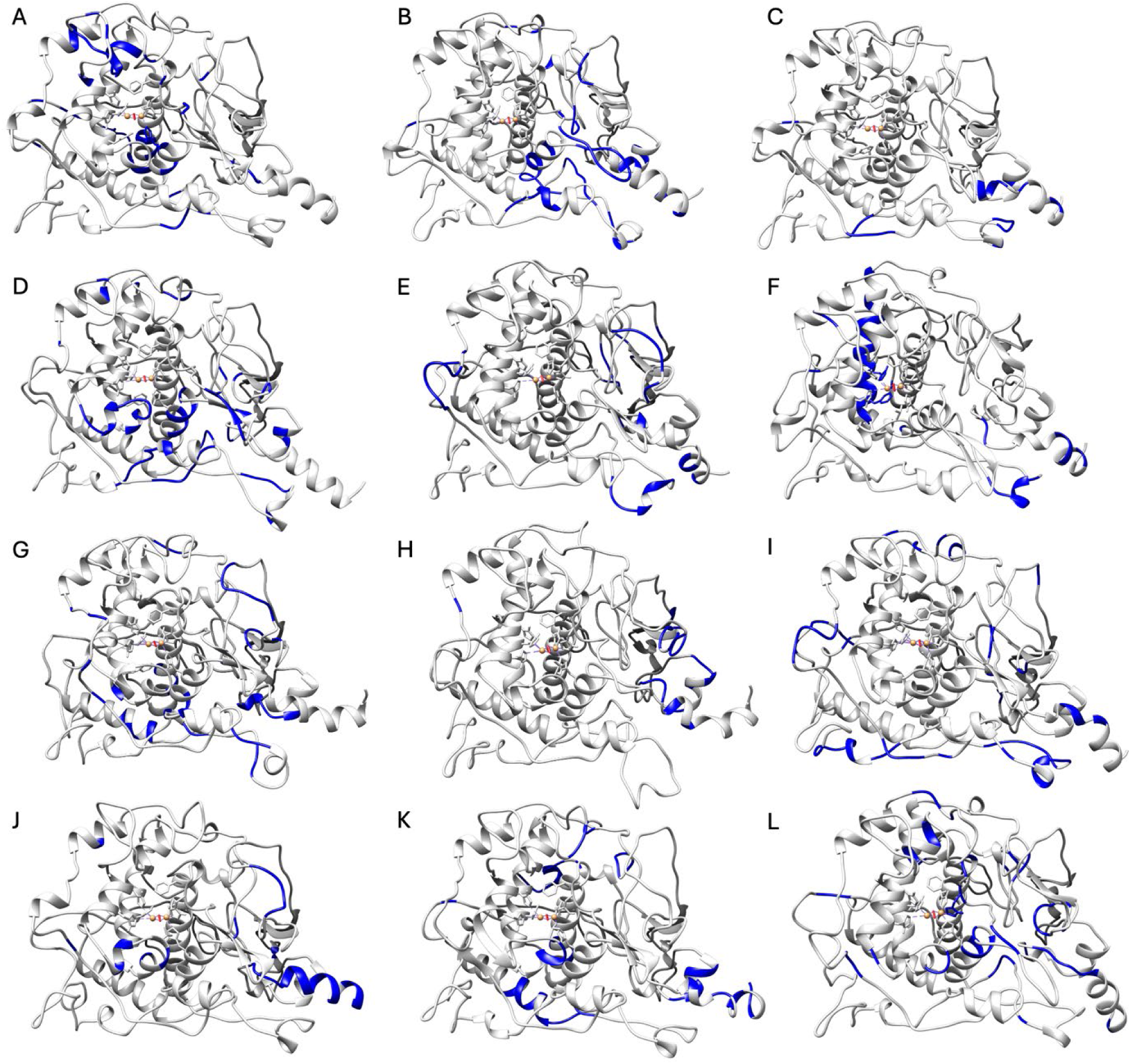
Top 10% loadings projected onto reference PDB structures. Visualization of top 10% of loadings (blue) for WT (Panels A-C, Clusters 1-3), R217Q (Panels D-F, Clusters 1-3), R402Q (Panels G-I, Clusters 1-3), and R217Q/R402Q (Panels J-L, Clusters 1-3) mutant variants are presented by cluster.

In the wild-type, most identified residues located within the alpha-helices making up the active site or in surrounding regions framing the active site. The exception is Cluster 3, which highlights residues further away from the active site. The negative PC1 subspace shared by Clusters 1 and 3 can be characterized by large loadings in the core alpha-helices H7, H9, and H10 (**Figure 1**) and at the junction of the C-terminus and loop. Cluster 2 can be represented by positive PC1 loadings, which include the beta-sheet residues M185-D199, H12 helix, and V150-I170 loop. Cluster 1 has unique characteristics as defined by positive PC2, corresponding with alpha-helices H15 (residues L340-D344) and H14 (residues F317-L322). Cluster 3 has unique characteristics as defined by negative PC2 in the signal peptide region and loops leading to H12 helix.

Residue analysis of each cluster’s top ten percent loadings provides insight into the variations in V150-I170 loop degradation in the mutant variant R217Q. In Cluster 1, H9 helix, portions of the beta-sheet residues M185-D199, and the region around the residue V337 constitute significant loadings. In Cluster 2, the alpha-helical region including residues V150-I170, several loop regions lateral to the active site appear characteristic. Finally, Cluster 3 also highlights the alpha-helical region (residues V150-I170) alongside the H19 helix. For the R402Q mutant variant, nearly all residues with large loadings for each cluster resided away from the active site and near the surface of the protein, most proximal to the N-terminus. Notably, Cluster 3 also contains the residues N300-P310. Differences between the three R402Q clusters and the WT protein are relatively minor.

The residues with the top ten percent loadings for the R217Q/R402Q mutant variant are quite variable across the clusters. The top loadings in Cluster 1 correlate primarily with the signal peptide and a nearby loop (P70-V74) and several residues proximal to residue V377 in the active site. The conformation of this loop may be the favorability between the Q68-D333 and E203/K334-D197 interactions. In this Cluster, this stems from the separation of residues D76, F84, and V83 (**Supplementary Figure S4**) Cluster 2 shares the residue V377 features with the addition of residues in helices H7 and H4 nearer the Cys-rich subdomain, and a sequence of residues located at the top of the active site, as pictured. Finally, Cluster 3 exhibits many residues concentrated at the junction of the Cys-rich and catalytic subdomain alongside a portion of the H9 helix.

### Tyr Coordinated Movement

Dynamical cross-correlation matrices (DCCM) display movement relationships between residues providing insights into large-scale protein motions (**Figure 5**).

**Figure 5.**
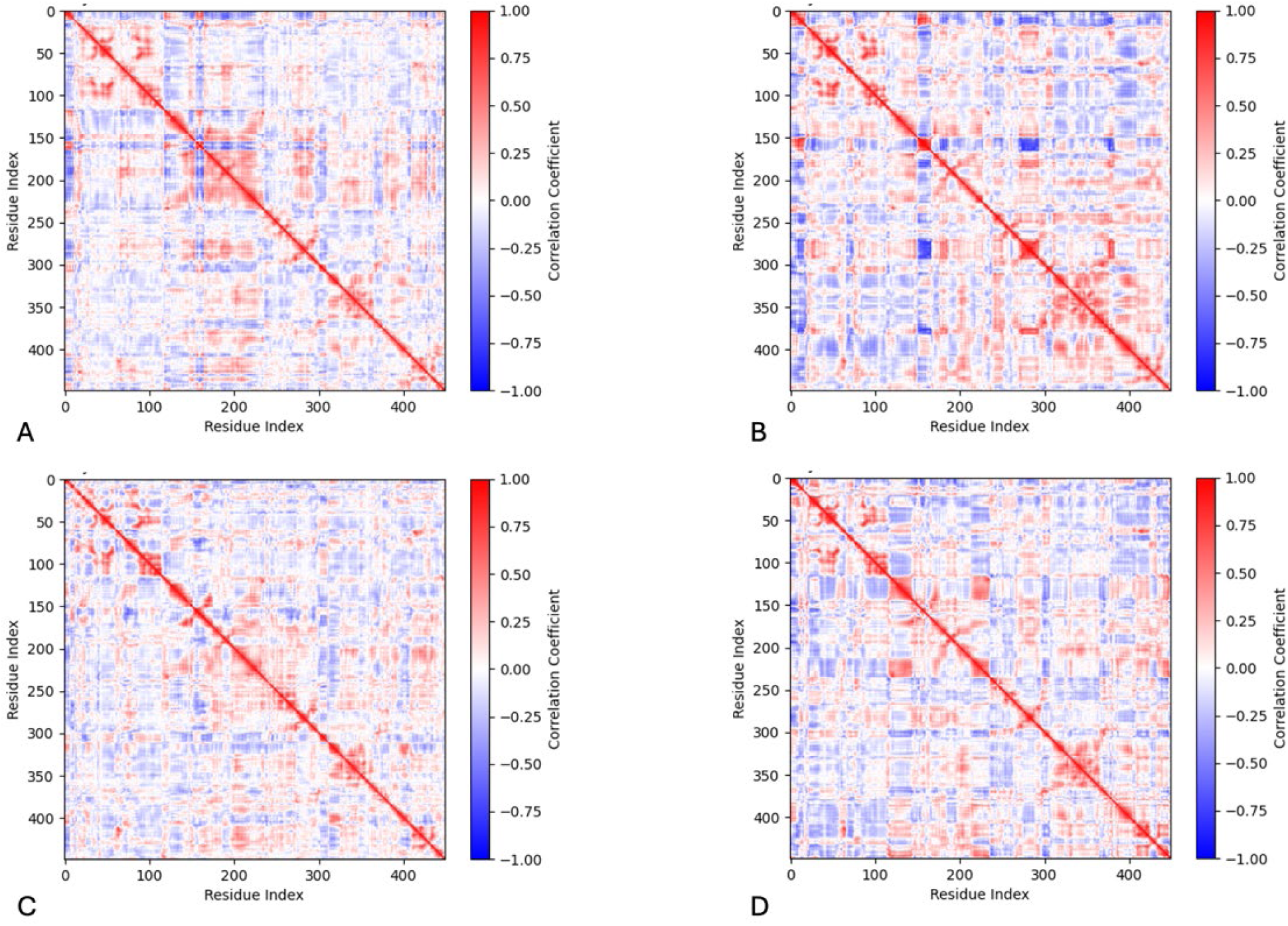
Dynamical cross-correlation matrices represent the movement relationships between amino acid residues. The matrices are shown for Tyr (Panel A), R217Q (Panel B), R402Q (Panel C), and R217Q/R402Q (Panel D) mutant variants.

The correlation coefficients range from +1, designating a strong positive correlation with red, to - 1, designating a strong negative correlation with blue. A correlation coefficient of 0 indicates no correlation and is colored white. There exist negatively correlated movements within the wild-type protein structure between residues Y156-P165 (helix H8) and M166-I233 (helices H9 & H10) (**Figure 1**). The regions which include residues M166-R196 (helix H9) and V150-T155 (helix H8) as well as M166-T226 (helices H9 & H10) with itself, move together. The latter coordinated movement is predominantly lost in each of the mutant variants. Residues M166-T226 (helices H9 & H10) is also positively correlated with R278-L297 (helix H12) and Y411-F429 (C-terminus). The R217Q mutant variant deviates from the wild-type pattern in many ways, with generally much stronger correlations. There is a stronger negatively correlated movement between the residues F269-P293 (helix H12) and residues P152-F167 (helix H8). The regions of F269-P293 (helix H12) and W178-L188 (helix H9) move together and so do the regions of Y411-I440 (C-terminus) and S267-N382 (helices H12-H16). The residues L263-I440 (helix H12 to C-terminus) and residues R239-P257 are positively correlated, and the R311-S315 region has a stronger positive correlation with itself. Finally, in the double mutant variant residues R116-T144 (helix H7) display coordinated movement with residues F214-W236 (helix H10), and there exists a smaller coordinate region around residue P310, like R217Q. Importantly, both R217Q and R217Q/R402Q mutant variants display negatively correlated movement between the residues at the junction of the catalytic and Cys-rich subdomains. The N-terminus region for the R217Q mutant variant includes residues L10-P21, S26-C112, and F124-I151, and the C-terminus region is residues D383-Q408. The N-terminus region for the double mutant variant is Y7-R115, and the C-terminus region is A381-S424 and R434-Y449. This coordination is stronger in the single mutant variant but spans larger regions of residues in the double mutant variant.

The coordinated movements can be visualized within a porcupine plot, where the magnitude and direction of movement for each alpha-carbon atom can be visualized with an arrow (**Figure 6, Supplementary Figure S5**). For this purpose, we used superimposed porcupine plots to show the differences in movements between mutant variant and wild type tyrosinase.

**Figure 6.**
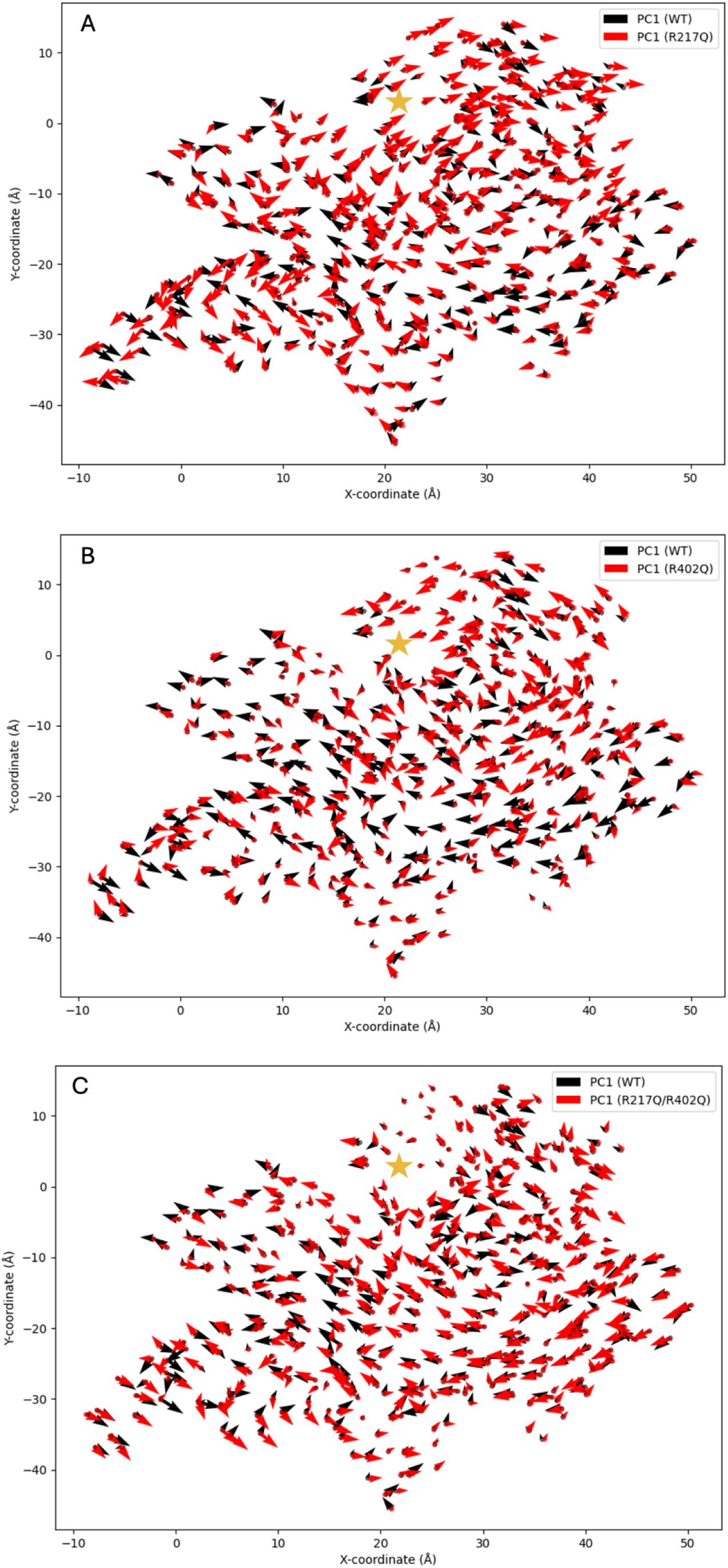
The superposition of porcupine plots of each mutant variant and wild type tyrosinase. Porcupine plots are shown for R217Q (Panel A), R402Q (Panel B), and R217Q/R402Q (Panel C) mutant variants. The approximate active site location is denoted with a gold star on Panel C.

Figure 7, Panel A displays the large increase in movement around the opening of the active site and in the core alpha helices within the R217Q mutant variant. In this mutant variant, decreased movement exists in the C100-L175 region (helices H7-H9). Shifts in the direction of movement can be viewed in the signal peptide, Cys-rich subdomain, and residues V150-I170 (helix H8). Panel B displays the decreased movement in R402Q within the region of residues L65-F200 (Cys-rich to helix H10). Increased movement is visible at the C-terminus and residues N300-F340 (H14) region at the active site. Changes in movement direction occur at the signal peptide, within the N300-F340 (helix H14) region, and some residues of the core alpha-helices. Finally, in Figure 7, panel C increased movement can be visualized in residues R22-R52 (helix H4), Q90-I170 (Cys-rich to helix H9), C-terminus, and N230-D240. Shifts in the direction of movement between wild-type tyrosinase and the R217Q/R402Q mutant occur in the signal peptide, Cys-rich subdomain, as well as the residues L60-P70. Each variant displays unique changes in movement as compared to Tyr. Secondary shifts in movement, as defined by PC2, are also available with supplementary materials (**Supplementary Figure S6**).

**Figure 7.**
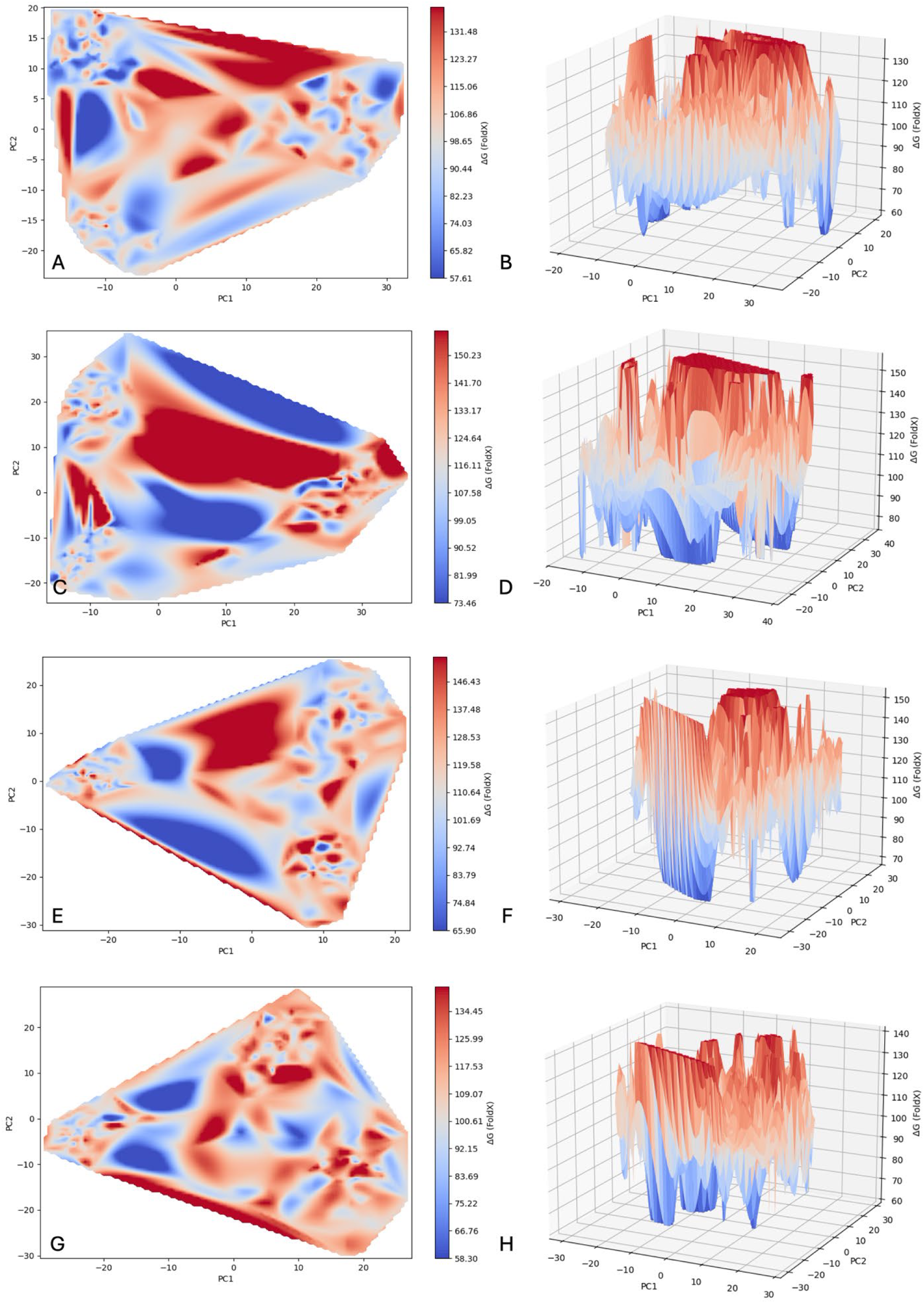
Tyrosinases Free Energy Landscapes. The landscapes are represented for wild type tyrosinase (Panels A, B), R217Q (Panels C, D), R402Q (Panels E, F), and R217Q/R402Q (Panels G, H). Mutant variants are shown in both 2D and 3D plots, respectively.

### Free Energy Profiles

Free energy profiles for wild type tyrosinase and the three mutant variants were calculated to understand the impact of structural and motion changes on protein stability. A free energy landscape for each of the four proteins was created by plotting Gibb’s free energy values for each timestamp in the concatenated trajectory against PC1 and PC2 (**Figure 7**).

Larger, positive values are indicated in red, smaller, positive values in blue, and values in between along this red-blue gradient. The presence of many peaks and valleys, giving a marble-like appearance, suggests that each protein is highly adaptable, taking many stable and unstable states within each cluster. Generally, the wild type tyrosinase has clusters at minima, though Cluster 2 displays higher ΔG values than the other two. Visually, it appears that a relatively low energy barrier exists between clusters 1 and 3, which share conformational characteristics with significant influence from residues in core helices. This suggests a stable, fluid form of Tyr in contrast with a less stable and unfavorable conformation with distinct properties. It shows a relatively low ΔG range as compared to each mutant variant (**Table 2**).

**Table 2.** Gibb’s Free Energy Descriptive Statistics.

| Structure | Average<br>(kJ/mol) | Minimum<br>(kJ/mol) | Maximum<br>(kJ/mol) | Trajectory 1<br>Average<br>(kJ/mol) | Trajectory 2<br>Average<br>(kJ/mol) | Trajectory 3<br>Average<br>(kJ/mol) |
| --- | --- | --- | --- | --- | --- | --- |
| Wild type | 97.0 ($\pm 13.6$) | 57.6 | 138.9 | 89.8 ($\pm 12.5$) | 103.4 ($\pm 13.8$) | 97.6 ($\pm 11.0$) |
| R217Q | 117.2 ($\pm 15.0$) | 73.5 | 157.9 | 125.8 ($\pm 15.2$) | 116.5 ($\pm 13.1$) | 109.4 ($\pm 12.0$) |
| R402Q | 112.7 ($\pm 14.2$) | 65.9 | 154.5 | 118.1 ($\pm 14.7$) | 112.9 ($\pm 14.1$) | 107.2 ($\pm 11.9$) |
| R217Q/R402Q | 109.3 ($\pm 13.0$) | 58.3 | 142.1 | 114.7 ($\pm 12.4$) | 103.6 ($\pm 10.8$) | 109.6 ($\pm 13.4$) |

R217Q has the highest minimum at 73.5 kJ/mol and maxima at 157.9 kJ/mol. Clusters 2 and 3 have lower average ΔG values than cluster 1, sharing a semi-stable conformation defined by partially coupled at residues150-170 and the signal peptide region. Unlike Tyr, all R217Q clusters have higher energy barriers, maintaining three distinct conformations and hindering conformational transitions. The R402Q mutant variant has a similarly high maximum to variant R217Q. Yet, total and individual trajectory averages remain lower than all corresponding R217Q values and higher than wild-type values. All clusters align with minima, and no structurally significant features align with stability patterns. The double mutant variant lies between the single mutant variants and the wild type, ranging from about 60-140 kJ/mol. Clusters 1 and 3 share higher average ΔG values and defining characteristics external to the core helices. These clusters appear to align with peaks, or relative maxima, likely indicating low mobility between conformational energy states. This contrasts with the more stable conformation of cluster 2, which derives its distinct conformation with significant influence from core alpha helices. Generally, Tyr displays more conformational mobility and stability as compared to variants.

In addition to the free energy landscapes, ΔΔG values for each mutant variant were calculated using a combination of the Internal Control Python code created for use with the Unfolding Mutation Screen (UMS) and FoldX. With these tools, several methods of predicting ΔΔG stability values for double mutation variants were tested alongside the use of DDGun (**Supplementary Table S4)**. For each method, the input structure, time under MD, ΔΔG, and P-value before UMS were recorded. The best method, determined by the mutant lowest structural P-value, used the input of an already mutated R217Q tyrosinase structure. The mutation was completed in Yasara. This structure underwent 2 ns MDs before reaching an acceptable P-value of 4.14E-10. Then the structure was input into UMS where the ΔΔG value assigned to R402Q was extracted. This stepwise method of mutating the structure produced a ΔΔG value of 1.73 kJ/mol, very similar in magnitude to the other FoldX methods, providing more realistic ΔΔG values when compared to the DDGun, another program capable of calculating ΔΔG for multiple mutation variants.

### Flexible Junction of Cys-rich and Catalytic Subdomains as Potential Location for Tyr Instability

The correlated movements and stability analyses of the four protein structures studied here provide insights into the flexible regions of Tyr (**Figure 8**). The R217Q mutant variant displays this instability through visible changes of the protein’s surface, at least in part caused by a reduction in stabilizing contacts with W178, where large-scale separation of the 150-170 (H8) loop and signal peptide from the M185-D199 beta sheets occur (**Supplementary Figure S7, Supplementary Table S5**). MoleOnline was used to analyze the presence of tunnels within the wild-type and R217Q/R402Q mutant variant structures. A dynamic, omnipresent network of tunnels proximal to H9 and H10 is characteristic of the wild-type structure, providing support for this portion of the tyrosinase protein being particularly susceptible to mutation-driven instability.

**Figure 8.**
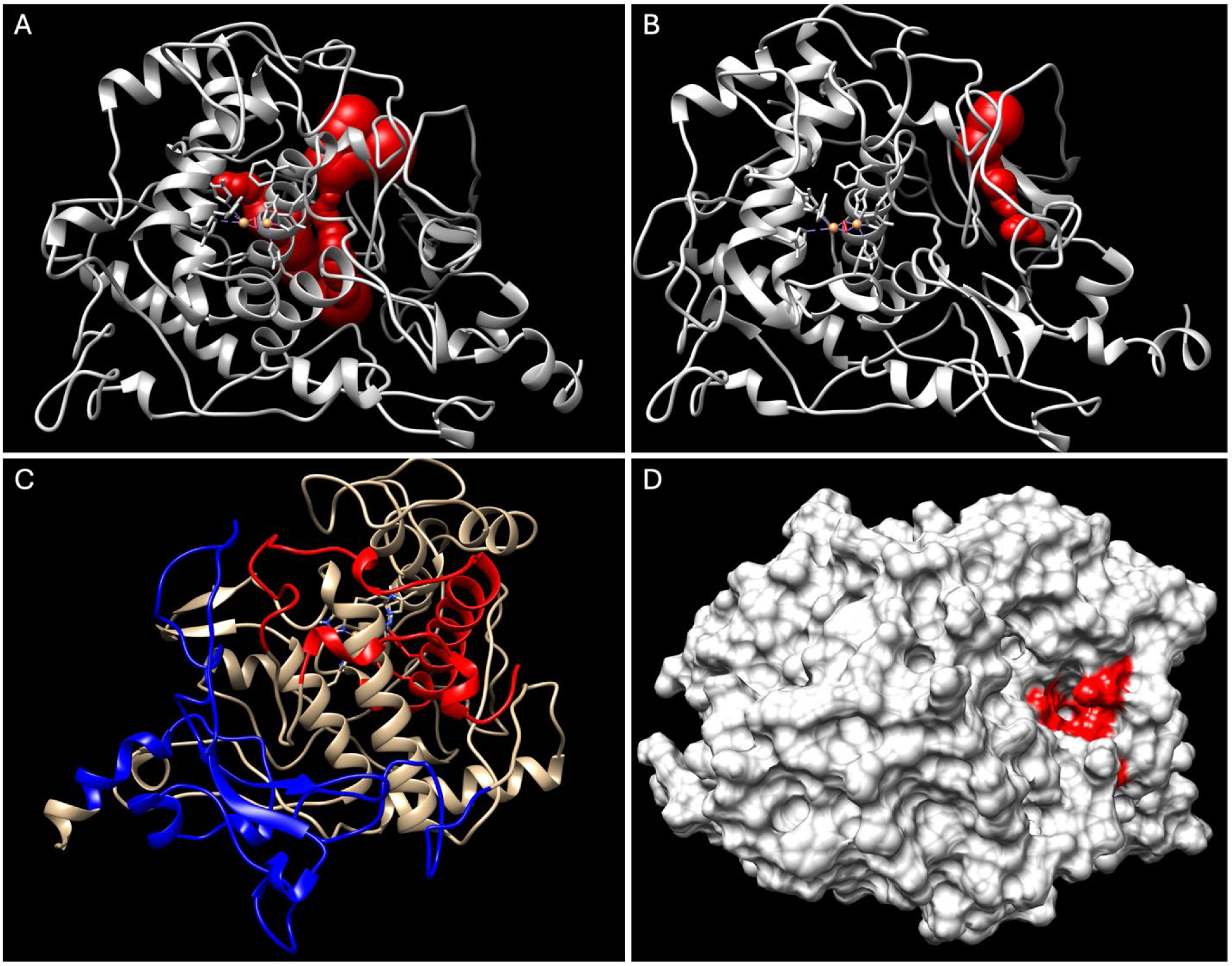
Tunnel formation correlates with flexible Tyr regions. Tunnels are shown in wild-type Tyr (Panel A) and R217Q/R402Q mutant variant (Panel B). The N-terminus (residues Y7-R115) and C-terminus (residues A381-S424, R434-Y449), between which the tunnel forms in the R217Q/R402Q mutant variant are shown in red and blue, respectively (Panel C). The surface rendition of the tunnel opening shows residues lining the tunnel (Panel D, red).

Interestingly, the R217Q/R402Q mutant variant exhibited a higher frequency of an additional tunnel between the Cys-rich and catalytic subdomains (**Table 3**).

**Table 3.** Two-proportion Z-test for tunnel analysis of wild-type and R217Q/R402Q structures.

|  | WT | R217Q/R402Q |
| --- | --- | --- |
| **Frames with Tunnel** | 23 | 38 |
| **Total Frames** | 60 | 60 |
| **Proportion** | 0.38 | 0.63 |
| **Combined Proportion** | 0.51 |  |
| **Standard Error** | 0.09 |  |
| **Z-Score** | -274 |  |
| **P-Value** | 0.0062 |  |

Though present in the wild-type structure as well, a two-proportion Z-test generating a P-value of 0.0062 suggests a significant difference in the presence of this tunnel system between wildtype and mutant tyrosinase protein structures. Both wild-type and double mutation variants were sampled every 5 ns from all three trajectories, resulting in a 60 timeframe samples. The segment of residues lining this tunnel from both subdomains aligns with the residues Y7-R115 and A381-S424, R434-Y449, identified within the dynamic analysis, suggesting that the presence of this tunnel, or the increased liminal space it represents, is the product of these molecular movements.

## Discussion

Dimensional reduction techniques, such as PCA, and free energy profiles are crucial in the systematic and comprehensive understanding of protein dynamics.^31^ Neither of these has previously been applied to the study of tyrosinase proteomics, neglecting large-scale conformation dynamics. Therefore, in this study, PCA was utilized jointly with dynamic analyses and free energy profiles to describe the coordinated movements of tyrosinase for the first time alongside selected mutant variants – R217Q, R402Q, R217Q/R402Q. While our previous study explored the underlying mechanism of instability for each of these variants, the current exploration advances our understanding of the coordinate movements fundamental to Tyr stability and their susceptibility to disruption^29^. We show that the PCA-identified coordinated movement underlying the stable conformations of wild-type tyrosinase arises within H9 and H10 helices, which is subsequently lost in disease-causing R217Q and R217Q/R402Q mutant variants. These helices are proximal to the large, flexible tunnel system and the interface of the catalytic and Cys-rich subdomains. We also characterize coordinated movements and energy profiles for each mutant variant to demonstrate that this flexible interface is a likely site for mutations to induce instability. Therefore, we suggest the utilization of chemical chaperones for the stability of this region, describing potential binding pockets for the small, hydrophobic Afegostat.

### Molecular Dynamics

We first conducted 100 ns molecular dynamic simulations for each of the four protein structures. The RMSF-colored PDB structures highlight the relatively low movement of each protein core. It also displays notable movements not previously explored, including increased RMSF in the residues N300-L312 region and proximal to the N-terminus – hairpin turn of beta sheets (residues A187-W195) and residues C35-G41 – which present elevated as compared to wild-type Tyr. The RMSF analysis reveals elevated movement in regions along the interface of the Cys-rich and catalytic subdomains and near the N-terminus as characteristic across variants. The SASA method does not meaningfully differ between tyrosinase proteins, instead displaying internal variation between clusters. Similarly, Rg shows a lack of distinction between proteins and similar variation amongst clusters. The SASA and Rg findings predominantly suggest that differences in protein stability between Tyr and variants are not attributable to complete protein unpacking, instead a nuanced understanding of instability. However, differences between clusters can provide insights into the flexibility and diverse conformations assumed by each Tyr protein.

### PCA

The essential dynamics of each protein structure was analyzed using PCA. using. The analysis showed that snapshots for each protein form clusters by trajectory with mild overlap, sharing prominent features. In each case, two clusters share half the PC1 subspace differentiated by a sign in the PC2 subspace. For Tyr, the similarities of the two clusters can be characterized according to high loadings in H7, H9, and H10 – core alpha-helices connected to residues 150-170 loop and lining the interface of the subdomains – with more nuanced differences more distal to the protein core. These differences in smaller helices framing the outer boundary of the active site may account for variable active site conformation and hold catalytic relevance in the determination of substrate binding and specificity. The dissimilar cluster exhibits prominent features along the outer perimeter of the active site and adjacent to the N-terminus. A notable difference in the R217Q mutant variant is prominently featuring minimal portions of helices H7 and H10 and larger portions of helix H9 with oppositely charged loadings to the residues 150-170 loop and signal peptide. This suggests a lack of coordinated movement in H7 and H10 helices and opposing movements of the two protein units, as compared to Tyr. The R402Q mutant variant displays reduced loadings in core alpha-helices, primarily in the H7 helix, exhibiting prominent features peripherally. Interestingly, the R217Q/R402Q mutant variant exhibits defining characteristics within H7, H19, H10, and small portions of H9 helices. Each variant alters the purpose and degree of influence of the core helices on the overall Tyr protein structure.

### Tyr Coordinated Movements

Next, a combination of DCCM and porcupine plots was used to describe the coordinate movements within each protein, building upon the structural classifications defined by PCA. Tyr displays a large, positively coordinated region encompassing H9 and H10 helical structures. It is plausible that this coordinated movement within the core helices is responsible for maintaining proper active site conformation while preserving overall structure flexibility. This region, well defined in Tyr, is lost to varying degrees in each mutant variant, displaying lower correlation values and disordered directionality. As noted in the prior paragraph, the R217Q/R402Q mutant variant has important features within the helical bundle like Tyr. However, combining this information with a dynamic understanding shows that in the case of the double mutation, these high loadings are indicative of uncoordinated, likely disruptive motions rather than the stabilizing effect of the wild type. Thus, each mutation may impact the integrity of the catalytic site as well as the structural stability. Interestingly, the R217Q and R217Q/R402Q mutant variants display an added region with negatively coordinated movement not seen in the wild type or R402Q mutant variant. These regions are the N- and C-termini, which line the interface of the Cys-rich and catalytic subdomains. The fundamental motions within the Tyr helical bundle play a large role in stabilizing the protein tertiary structure and catalytic site. When disrupted through mutation, the lack of coordinate movement may lead to reduced stability in all areas of the protein, primarily at the junction of the two subdomains.

### Free Energy Profiles

Gibb’s free energy landscapes were obtained for each mutant variant to determine the effect of the structural and dynamic differences on Tyr stability. All four proteins display a range of available conformations, with each mutant variant maintaining higher ΔG values than the wildtype. Notably, the two clusters sharing features with significant influence from helices H7, H9, and H10 display lower average ΔG values providing evidence that the coordinated movement by these core helices imparts stability to the Tyr protein. Thus, this coordinated movement, when lost as it is presumably in the third Tyr cluster as well as the variants, results in a less stable protein structure. Wild type Tyr also appears to have a lower energy barrier between the two similar clusters than with the third, less stable cluster or between the clusters in the other protein landscapes. This suggests the wild type protein has a higher degree of conformational mobility and structural stability than each mutant variant.

### Tyr Instability at Interface of Cys-rich and Catalytic Subdomains

Finally, an analysis describing tunnel systems proximal to the interface of the Cys-rich and catalytic subdomains in the wild-type and R217Q/R402Q mutant variants provides additional evidence for the region flexibility and concurrent structural liability. Tyr naturally contains an extensive, mobile tunnel system roughly located at this interface. One of this tunnel system’s associated surface cavities has been studied previously, showing a negative correlation between cavity size and OCA1B mutant protein stability (ΔΔG)^9^. It follows that the mutation characteristics observed in this paper – larger cavity volume and SASA – are underlying fundamental changes described in this study. As in the case of the R217Q/R402Q mutant variant, this increased cavity size can be understood in terms of an additional tunnel forming separation between the two subdomains. The flexibility of the Tyr protein to maintain this tunnel system can therefore be understood not only as a structural adaptability advantage but also as a mutation susceptibility disadvantage.

### Western blot

Interestingly, performing a Western blot on the protein lysates may provide experimental insight into the shared instability of the R217Q and R217Q/R402Q mutant variants. The lysates for all four truncated tyrosinase proteins were prepared and purified as previously described^29^. Western blots for the lysates were completed using three different primary antibodies: Santa Cruz Biotechnology mouse monoclonal (T311), Santa Cruz Biotechnology rabbit polyclonal with epitope 421-529 (H-109), Invitrogen mouse monoclonal (C2-85 binds the epitope 51-100) (**Supplementary Figure S8**). All Western blots exhibit a Tyr band for each protein around 68 kDa. The R217Q and R217Q/R402Q variants also display a smaller, distinct band around 54 kDa. The WT and R402Q also display this band, however, it is very faint and difficult to tell if the stain is just a byproduct. The difference of approximately 14 kDa equates to a roughly 127 residue loss to produce this smaller band. To determine if the protein sequence had truncations at either terminus two antibodies were chosen. The H-109 antibody binds the epitope in positions from 421-529, determining the presence of the C-terminus within the additional smaller band.

The C2-85 antibody binds amino acids in positions 51-100, determining the presence of the N-terminus in this second band. In both instances the smaller band persists, suggesting that amino acid residues may be proteolytically digested at both termini. This aligns with the computational results of the R217Q and R217Q/R402Q mutant variants, which demonstrate increased dynamics at both termini lining the junction of the Cys-rich and catalytic subdomains.

### Potential Chaperone Protection of Tyr-Coordinated Movements

Thus, we decided to explore the theoretic application of chemical chaperone to reinforce this area of tyrosinase. Previous work has demonstrated the successful use of chaperone molecules in treating various diseases^24; 27; 28^. A recent publication has shown the rescue of Tyr activity and melanin production using chemical chaperones 4-phenylbutyric acid (4PBA) and tauroursodeoxycholic acid (TUDCA)^48^. We expect that the flexible portions of the tyrosinase structure, primarily at the junction of the Cys-rich and catalytic subdomains, may be stabilized by such chaperone molecules. Specifically, we docked a known chaperone molecule, Afegostat, to the wild type tyrosinase structure to identify potential chaperone binding locations. (**Figure 9**).

**Figure 9.**
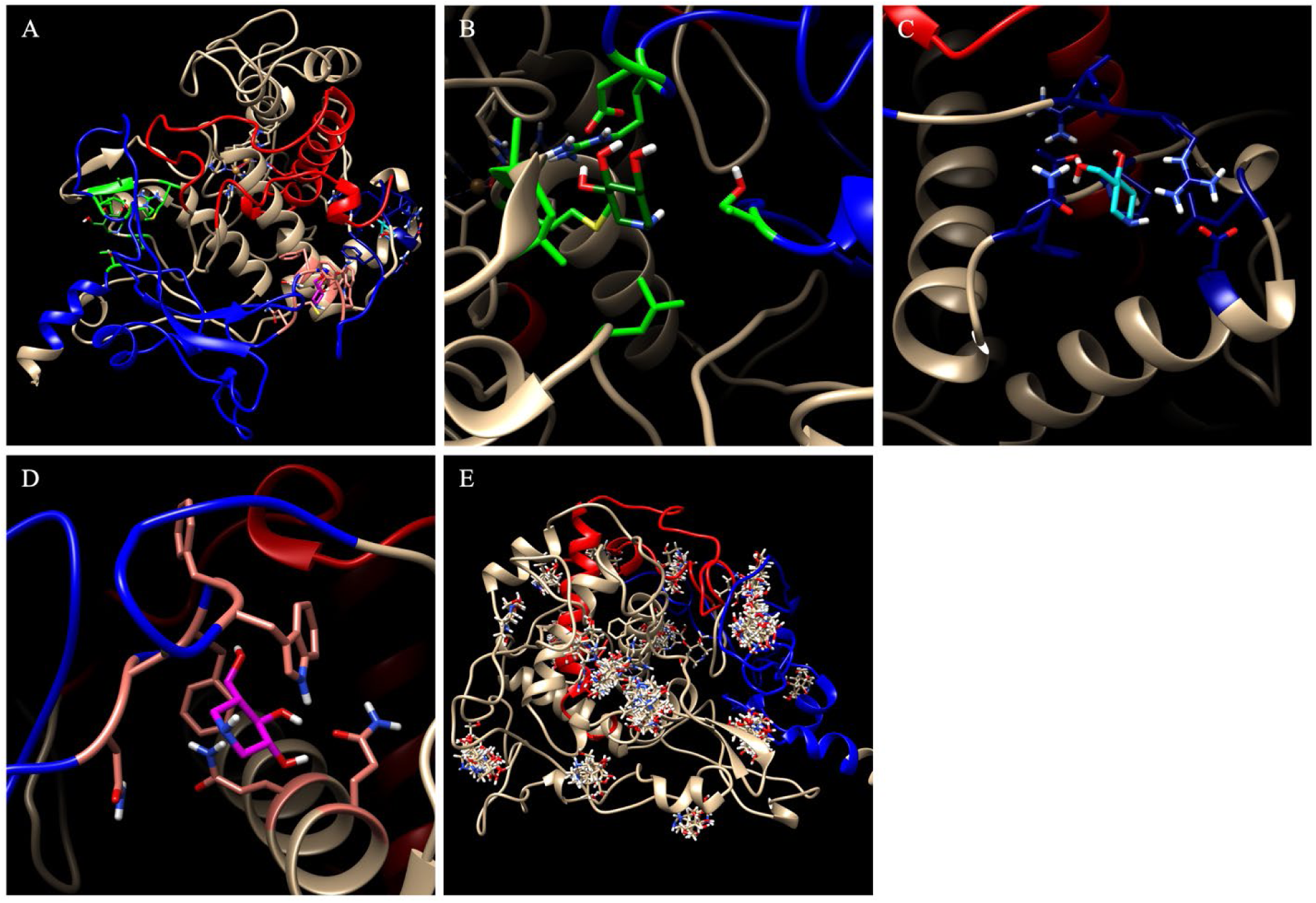
Potential tyrosinase binding pockets for chaperone molecules. Shifting regions containing residues Y7-R115 and A381-S424, R434-Y449 are shown in blue and red, respectively. Three potential binding pockets (Panel A) and Afegostat are highlighted in light/dark green (Panel B), dark/light blue (Panel C), and pink/magenta (Panel D). All potential clusters are also shown (Panel D).

The top three candidates are shown proximal to this interface (**Figure 9A**). The first exists at the tetrameric junction of the signal peptide, residues V150-I170 loop, beta sheets, and residues L279-S284 alpha-helix (**Figure 9B**). The docking of Afegostat may stabilize this tetrameric junction, increasing stability between the two tyrosinase subdomains. The binding energy for this position is 4.986 kcal/mol. The second potential binding site exists at the back of the active site (**Figure 9C**). Here, afegostat can stabilize both helices H7 and indirectly H10. These helices are involved in stabilizing the V150-I170 loop which experiences instability in each of these mutant variants. The binding energy for this position is 5.175 kcal/mol. The third binding site exists between the loop region succeeding the Cys-rich subdomain and H10 helix, directly binding to residues of both other sites (**Figure 9D**). In this location, Afegostat may directly stabilize the junction of the subdomains. The binding energy for this position is 4.975 kcal/mol. Additionally of the 100 docking poses produced by Yasara, 59 poses bound to at least one of these two regions accounting for 12 of the 20 total clusters (**Figure 9E**). Each of these locations may be prospective mechanism to use small, hydrophobic chemical chaperones like Afegostat to stabilize tyrosinase mutations impacting the stability of the Cys-rich and catalytic interface. Enhancing the stability of these mutants through chemical chaperones may provide a therapeutic approach to recover tyrosinase activity in patients suffering from OCA1-causing genetic mutations, such as the R217Q and R217Q/R402Q mutant variants.

### Signal Peptide Importance in Understanding Tyr Endoplasmic Reticulum Retention

Notably, the molecular dynamics completed in this paper use a model, which includes the signal peptide region of the Tyr protein. This is important because the signal peptide while integral to Tyr transport, is believed to be cleaved and thus not present intramelanosomally^49; 50^. Thus, our representation with the signal peptide provides insights into Tyr folding and stability pre-trafficking to the melanosome. This is particularly relevant to Tyr, as mutant variants leading to OCA1 have been shown to have increased endoplasmic reticulum (ER) retention, slowing or inhibiting protein transport to melanosomes^51-53^. Various chemical chaperones, including substrates L-tyrosine and L-DOPA, have also been studied in vitro with promising results to aid in proper tyrosinase folding within the ER, improving both Tyr catalysis and melanin production^25; 48^. In this study, we show destabilization at the junction of the Cys-rich and catalytic subdomains which is proximal to, and in some cases directly impacting the movement of the signal peptide. We have also provided corresponding potential locations for chemical chaperone binding. Thus, we speculate that chemical chaperones improving Tyr protein folding within the ER may play a role in stabilizing the interface of Tyr subdomains and this signal peptide.

### Limitations & Future Directions

The rationale behind PCA, which makes it a useful tool– dimensionality reduction– is also its limitation. Dimension reduction techniques remove data, particularly data relating to more nuanced movements, in favor of simplicity. While this allows for a broad, statistical approach to protein dynamics, capturing only ‘essential’ movements, it cannot provide a more detailed analysis. Maximizing the total variance captured in the first few principal components is quite challenging in large systems. Thus, it may be prudent to study individual regions of the tyrosinase structure with PCA to supplement the essential dynamics in this paper. These regions may include individual subdomains, the active site, and other targeted regions. This may provide more specific information on biologically relevant motions, especially when combined with other modeling techniques (docking) and experimental methods.

In addition, another limitation arises in our ability to comprehensively explore the energy landscape for tyrosinase and variants. While concatenation of triplicate trajectories allows for an increased sampling of tyrosinase subspaces as opposed to a single trajectory, it lacks the completeness that may be attained by collective variable-based sampling, such as Umbrella Sampling (US)^54; 55^. These methods track a few key variables over, allowing for a larger sampling of a smaller system, and the US, specifically, applies constraints to allow for sampling of rarer, high-energy states that may not be achieved in traditional methods^54; 55^. However, the trade-off remains that the US requires higher computational power than traditional molecular dynamics and cannot be completed for large, whole protein systems. Therefore, given technological and resource constraints, both methods used in tandem provide the best approach to contemporary dynamic and energy modeling.

Further steps include applying experimental methodologies, such as mass spectrometry, to determine the exact identity of the excised regions in the R217Q and R217Q/R402Q mutant proteins, and whether and to what extent this cleavage occurs in tyrosinase and other mutant proteins. *In silico* investigations should also be completed exploring the best-suited classes of molecules and potential mechanisms for tyrosinase stabilization with chemical chaperones. Subsequently, these might be tested *in vitro* to develop pharmacologic chaperones and therapeutics to recover tyrosinase activities in OCA1 mutant variants.

### Conclusions

Our multi-faceted analysis of tyrosinase and selected mutant variants is filling the gap in the absence of literature exploring the large-scale coordinated movements of the tyrosinase protein and broader category of melanogenesis proteins. With this, we demonstrated the flexibility of the junction between the Cys-rich and catalytic subdomains and its congruent susceptibility to instability. We also have provided the first free energy profile for tyrosinase, demonstrating several stable and high-energy, transition protein states, each with defining characteristics. Finally, we also calculate the change in Gibb’s free energy for a tyrosinase multiple mutation variant, the double mutation variant R217Q/R402Q. Succinctly, our study provides the basis for further tyrosinase dynamics and docking studies, exploring the therapeutic possibilities of chemical chaperones for OCA1 patients.

## Methods

### Lysate Preparation & Western Blots

The human recombinant intra-melanosomal domain of tyrosinase and mutant variants R217Q, R402Q, and R217Q/R402Q were expressed in *T. ni* larvae (C-PERL & Allotropic Tech, MD) [Dolinska 2014]. The larval biomass was homogenized in lysate buffer (20 mM sodium phosphate, pH 7.4, 500 mM NaCl, 5 mM imidazole, 2 mM MgCl2, 25 uL 1-phenyl-2thiourea (Sigma-Aldrich), 40 ug/mL DNAse I (Thermo Fischer), 0.2 mg/mL chicken lysozyme (Sigma-Aldrich), and protease inhibitors (Roche)). The lysate was incubated while rotating at 25 °C for 30 minutes, sonicated at 25 °C, and then centrifuged at 4 °C for 30 minutes at 8,000 RPM. The supernatant was then diluted 1:1 with affinity binding buffer (20 mM sodium phosphate, pH 7.4, 500 mM NaCl, 20 mM imidazole). All lysates were stored at -20°C.

The presence of wild type tyrosinase and each mutant variant in the lysate was observed using Western blot analysis using several anti-tyrosinase primary and anti-mouse secondary antibodies. First, the Santa Cruz Biotechnology T311 monoclonal mouse IgG2a tyrosinase antibody was used for its high target specificity and sensitivity. Next, the Santa Cruz Biotechnology H-109 antibody was used for its specificity in binding the epitope 421-529. Finally, the Thermo Fisher Scientific Invitrogen C2-85 tyrosinase monoclonal antibody was used for its specificity in binding the epitope 51-100. All proteins were run at a concentration of 0.5 mg/mL. The protein concentration of each variant was monitored at each step using the 12-channel microvolume spectrophotometer N120 (IMPLEN, VA, USA).

### Homology Model & Channel Visualization

A homology model of the intra-melanosomal domain of human tyrosinase (residue 1-449) was modeled using the NEI Commons Ocular Proteomes Tyrp1 atomic model (PDB:5M8L, https://neicommons.nei.nih.gov/#/proteomeData). Three mutant variants were generated from this model using the Edit > Swap > Residue function in Yasara: R217Q, R402Q, and R217Q/R402Q. The structures were energy minimized using the Options > Choose experiment > Energy minimization function in Yasara (http://www.yasara.org).

Alignments and visualizations for all five variants and the original tyrosinase structures were generated using UCSF Chimera, version 1.17.3 (https://www.cgl.ucsf.edu/chimera/). The superposition of the mutant variant and the wild type tyrosinase models after 100 ns of MD simulation was completed using the Tools > Structure Comparison > Matchmaker tool. The individual hydrogen bonds were generated using the Tools > Structure Analysis > FindHBond tool. Hydrophobic contacts were generated using the Tools > Structure Analysis > Find Clashes/Contacts tool. Additionally, the secondary structure and topology diagrams intended for the use of consistent secondary structure naming were created for the wild-type tyrosinase protein using the PDBSum Generate server. The input structure was the energy-minimized homology model previously described.

MOLE online (https://mole.upol.cz/) was used to generate tunnels within the wild-type and double-mutation variant models every 5 ns of the concatenated trajectories, a total of 60 structures for each protein. Channel parameters were set to an origin radius of 10 Å, a surface cover radius of 10 Å, and a Voronoi scale weight function, with the merge pores and automatic pores functions turned off. All other parameters and functions were standard. Tunnel visualizations were downloaded from the online server and viewed in Chimera. The number of time frames with and without a tunnel system between the Cys-rich and catalytic subdomains was computed for both the wild-type tyrosinase and the R217Q/R402Q mutation variant. A two-proportion Z-test was completed to determine if the observations between the wild-type and mutant were statistically significant.

### Root Mean Square Fluctuation, Solvent Accessible Surface Area, & Radius of Gyration Calculations

For wild-type protein and all mutant variants calculated in triplicates, root mean square fluctuation (RMSF) for each trajectory was calculated using the visual molecular dynamics program, VMD (https://www.ks.uiuc.edu/Research/vmd/). For use in VMD, the sim files were transferred to an xtc file using the Yasara ‘md_convert.mcr’ macro. The energy-minimized protein structure from each trajectory was chosen as the reference structure. The xtc file from Yasara was uploaded into this reference pdb. The snapshots were aligned first based on Cα atom position using the Trajectory > Align tool. After aligning the snapshots with the reference structure, RMSF was calculated by running a TCL file in VMD’s TK Console. The RMSF values were combined to reflect the final concatenated trajectory. Both solvent-accessible surface area (SASA) and radius of gyration (Rg) were calculated for each protein’s triplicate data using the ‘md_analyze.mcr’ macro in Yasara. Both sets of values, respectively, were combined to reflect the final concatenated trajectory of each protein structure.

### Molecular Dynamics & Principal Component Analysis

Like that previously described^29^ Each structure underwent a 100 ns MD simulation (MD_run.mcr) with snapshots generated every 1 ns, and this was completed for each in triplicate. The simulation conditions were as follows: AMBER14 force field, temperature of 298 K, pH of 7.4, water density of 0.997 g/mL, NaCl concentration of 0.9%, and a cubic cell extending 10 Å beyond the protein (94.6 Å × 94.6 Å × 94.6 Å). Initial atomic velocities were varied by altering the Randomized Seed for each simulation of the triplicates.

We completed principal component analysis for the concatenated trajectories of each protein as described.^37^. The molecular dynamics trajectories were converted from a series of sim files to xtc format for all proteins. The converted trajectories for each protein were concatenated, and the atomic positions were aligned using a Python script with the help of CoPilot (https://copilot.microsoft.com/). The ‘align’ function within the MDAnalysis package (https://www.mdanalysis.org/) was used for the alignment of the concatenated trajectory to a reference structure. Next, the atomic positions for the alpha carbon atoms were extracted using an additional Python script. The coordinate data was input into GraphPad Prism (https://www.graphpad.com/features) in a format where the columns represented the x, y, or z coordinate for each residue for a total of 1347 columns and where the rows represented each time frame for a total of 303 rows (including the 0 ns time frame).

The Prism PCA tool was utilized to standardize the data and then apply a parallel analysis to the previously described data frame using standard settings and according to Prism’s online manual (https://www.graphpad.com/guides/prism/latest/statistics/stat_pca.htm). Score and loading values were extracted to generate score and loading plots for each protein. To determine collections of timeframes, a silhouette analysis was applied to the score data using a K-means method of clustering56. The top-loading values were extracted for further simplicity in the analysis. Two sub-methods were employed to determine the cutoff for the highest loading magnitudes. The first method explored the full PC1 and PC2 subspaces, setting the threshold at |x| ≥ 0.7, where x is the loading value for any given variable. The second method separated loadings by cluster, assigning each to a cluster using the K-means Clustering and setting the threshold to include only the top 10% of loading values. In this second method, the magnitude of the loading was determined by Euclidian distance from the origin.

### Dynamical Cross-Correlation Matrix & Porcupine Dynamic Analysis

A dynamic cross-correlation matrix (DCCM) plot was created using a Python script,^38; 57; 58^ and the MDAnalysis package to align the concatenated trajectory to a reference structure. Then extracted coordinates of alpha carbon atoms were used to generate a correlation matrix using the NumPy package (https://numpy.org/). The correlation matrix was calculated by averaging dot products of the centered coordinates and normalized to a range of -1 to +1. Finally, the plot was created using Matplotlib (https://matplotlib.org/). Using the same packages, a 2D porcupine plot^38^ was also created with Matplotlib. Precomputed principal component eigenvectors were used from the Prism program, and the arrow representations symbolize both the direction and magnitude of each vector.

### Molecular Docking

A small chaperone molecule was docked to the tyrosinase protein surface to understand the role of molecular movements. A known chemical chaperone molecule was selected to perform docking to the wild type tyrosinase protein. The small, well-documented (3R,4R,5R)-5- (hydroxymethyl)piperidine-3,4-diol, also known as Afegostat, was chosen from PubChem (https://pubchem.ncbi.nlm.nih.gov/compound/447607) and docked to a 100-ns tyrosinase structure in Yasara using the ‘dock_run.mcr’ macro. The docking method VINA was used to complete a total number of 100 docking runs. All other parameters in the macro were left unmodified.

Potential chaperone binding sites were determined based on the following criterion. Given that Afegostat is hydrophobic, we presumed it to function as a hydrophobic rather than an osmolyte chaperone. Therefore, all clusters within the active site binding pocket were excluded as the hydrophobic chaperone molecules generally bind to exposed hydrophobic regions; the tyrosinase active site is primarily hydrophilic (Papp 2006; Cortex 2014). The potential binding pocket clusters were then sorted by binding energy and several members within the cluster. The cutoff for both parameters was set to the 75^th^ percentile of the given results. Therefore, the threshold for binding energy was 4.618 kcal/mol and a minimum of 7 docked Afegostat molecules per cluster.

### Gibbs Free Energy Landscapes & Change in Gibbs Free Energy

Gibbs free energy values were calculated for each 1ns timeframe for all concatenated trajectories using FoldX (https://foldxsuite.crg.eu/), using the RepairPDB command followed by the Stability command. A Python script was then created to generate 2D and 3D contour plots of the ΔG values within the PC1 and PC2 subspaces.

Change in Gibbs free energy was also calculated for the first time for the double mutant variant. The ΔΔG value was calculated using the FoldX-based Unfolding Mutation Screen (UMS).^59^. The input R217Q mutant variant PDB structure was energy minimized and input into the Internal control program to assess the P-value. The structure then underwent 2 ns of molecular dynamics before being reintroduced into the Internal control program, where it produced an acceptable P-value below 1×10^-6^. The BuildModel, RepairPDB, and Stability FoldX commands were applied to this structure. The R402Q ΔΔG value was then extracted from this stepwise method and assigned to the double mutation variant R217Q/R402Q. Finally, the DDGun online server (https://folding.biofold.org/ddgun/) and two other FoldX-based methods were also utilized as a comparison for the double mutation variant ΔΔG values (**Supplementary Table S4**).

### Supplementary Materials

The supplementary information includes additional descriptive statistics, silhouette analysis results, change in Gibb’s Free Energy (ΔΔG) values for R217Q/R402Q double mutation variant generated by various methods, PCA scree plots and loading analyses, labeled and PC2 porcupine plots, contacts with 3-dimensional surface renditions, and Western blots for tyrosinase and mutant variants.

## Supplementary Materials

**Paragraph S1.** Positive and negative PC1 and PC2 description

**Table S1.** Solvent Accessible Surface Area Descriptive Statistics

**Table S2.** Radius of Gyration Descriptive Statistics

**Table S3**. Silhouette analysis for tyrosinase and mutant variants.

**Table S4.** ΔΔG values generated by various methods.

**Table S5.** Residue W178 and V377 contacts in wild type tyrosinase and R217Q mutant variant..

**Figure S1.** Scree plots for wild-type tyrosinase and mutant variants are shown.

**Figure S2.** Representation of PC1 and PC2 subspace for tyrosinase and mutant variant ribbon structures.

**Figure S3.** The graphs obtained for the top 10% loadings per cluster are shown.

**Figure S4.** Instability in R217Q/R402Q mutant variants stems from the disruption of interactions.

**Figure S5.** PC1 porcupine plot for WT with residue identifiers.

**Figure S6.** The superimpositions of PC2 porcupine plots.

**Figure S7.** Instability in R217Q stems from the loss of interactions.

**Figure S8.** Western blots of tyrosinase and mutant variants.

## Acknowledgment

This research was supported by the Intramural Research Program of the National Eye Institute, ZIA EY000476-10 to Y.V.S.

Yuri V. Sergeev https://orcid.org/0000-0002-7204-6572

