## supplementary material for "Genetic mutations disrupt the coordinated mode of tyrosinase intra-melanosomal domain"

**Paragraph S1.** Positive and negative PC1 and PC2 description

For wild-type, the negative PC1 subspace represents residues in core alpha helices (H7, H9, & H10; **Figure 1**), the signal peptide, and the V150-I170 (helix H8) loop. The positive PC1 subspace overlaps with the negative PC1 subspace in the signal peptide and 150-170 loop regions while also including the beta-sheet residues M185-D199, a portion of the residues L279-S284 of H12 alpha helix, and loops between these regions and the active site. The negative PC2 subspace primarily represents residue loops along the protein's surface, while the positive PC2 subspace represents a few residues within the F317-L322 alpha helix (H14). In the R217Q mutant variant, the harmful PC1 subspace residues primarily exist within the signal peptide and V150-I170 loop. A few residues are also highlighted within core alpha helices. The positive PC1 subspace consists of many residues. Some notable regions include core alpha helices, the active site, and M185-D199 beta sheets. The negative PC2 subspace primarily highlights alpha helix 19, while the positive PC2 subspace includes surface-level loops. In the R402Q mutant variant, both the negative and positive PC1 subspaces are represented by residues within the external loops, alpha helices, and signal peptides. The same is true for both the negative and positive PC2 subspaces. Finally, the R217Q/R402Q mutant variant negative PC1 subspace includes many residues within the alpha helices 7, 10, and 19. The positive PC1 subspace includes external loops, alpha helices, as well as the M185-D199 beta sheets. The negative PC2 subspace consists of H15 and active site residues. The positive PC2 subspace is primarily constituted by the signal peptide.

**Table S1.** Solvent Accessible Surface Area Descriptive Statistics

| Structure | Average (Å$^2$) | Minimum (Å$^2$) | Maximum (Å$^2$) |
|---|---|---|---|
| Tyrosinase | 20633 (±380) | 19443 | 21282 |
| R217Q | 20884 (±458) | 19589 | 21941 |
| R402Q | 20974 (±337) | 19775 | 21802 |
| R217Q/R402Q | 20741 (±476) | 19674 | 21598 |

**Table S2.** Radius of Gyration Descriptive Statistics

| Structure | Average (nm) | Minimum (nm) | Maximum (nm) |
|---|---|---|---|
| Tyrosinase | 21.80 (±0.06) | 21.62 | 21.94 |
| R217Q | 21.91 (±0.21) | 22.49 | 22.41 |
| R402Q | 21.86 (±0.10) | 21.63 | 22.09 |
| R217Q/R402Q | 21.85 (±0.10) | 21.62 | 22.25 |

**Table S3**. Silhouette analysis for tyrosinase and mutant variants.

| Structure | Cluster Number | Silhouette Score |
|---|---|---|
| **Tyrosinase** | 3 | 0.7427 |
| **R217Q** | 3 | 0.7721 |
| **R402Q** | 3 | 0.7229 |
| **R217Q/R402Q** | 3 | 0.7044 |

**Table S4.** ΔΔG values generated by various methods.

| | | ΔΔG (kCal/mol) | | | | |
|---|---|---|---|---|---|---|
| *Method* | *Structure-MD Time* | *R217Q* | *R402Q* | *Double* | *P-value* | *Notes* |
| **Build Model Twice** | Emin-2ns | - | - | 1.44 | 4.69E-07 | |
| **Summation** | Emin-2ns | 0.343511 | 1.42263 | 1.77 | 4.69e-7 | R217Q \|\| R402Q |
| **Stepwise-R217Q** | Yasara-2ns | - | - | 1.73 | 4.14e-10 | |
| **DDGun** | 100ns MD | - | - | 0.4 | - | |
| **DDGun Summation** | 100ns MD | 0.6 | 0.1 | 0.7 | - | |

**Table S5.** Residue W178 and V377 contacts in wildtype tyrosinase and R217Q mutant variant.

| | Contacts | |
|---|---|---|
| **Structure** | **W178 with 150-170 \| 273-294** | **V377** |
| **Tyrosinase** | 9 \| 16 | 12 |
| **Cluster 1** | 4 \| 3 | 2 |
| **Cluster 2** | 11 \| 27 | 5 |
| **Cluster 3** | 5 \| 8 | 18 |

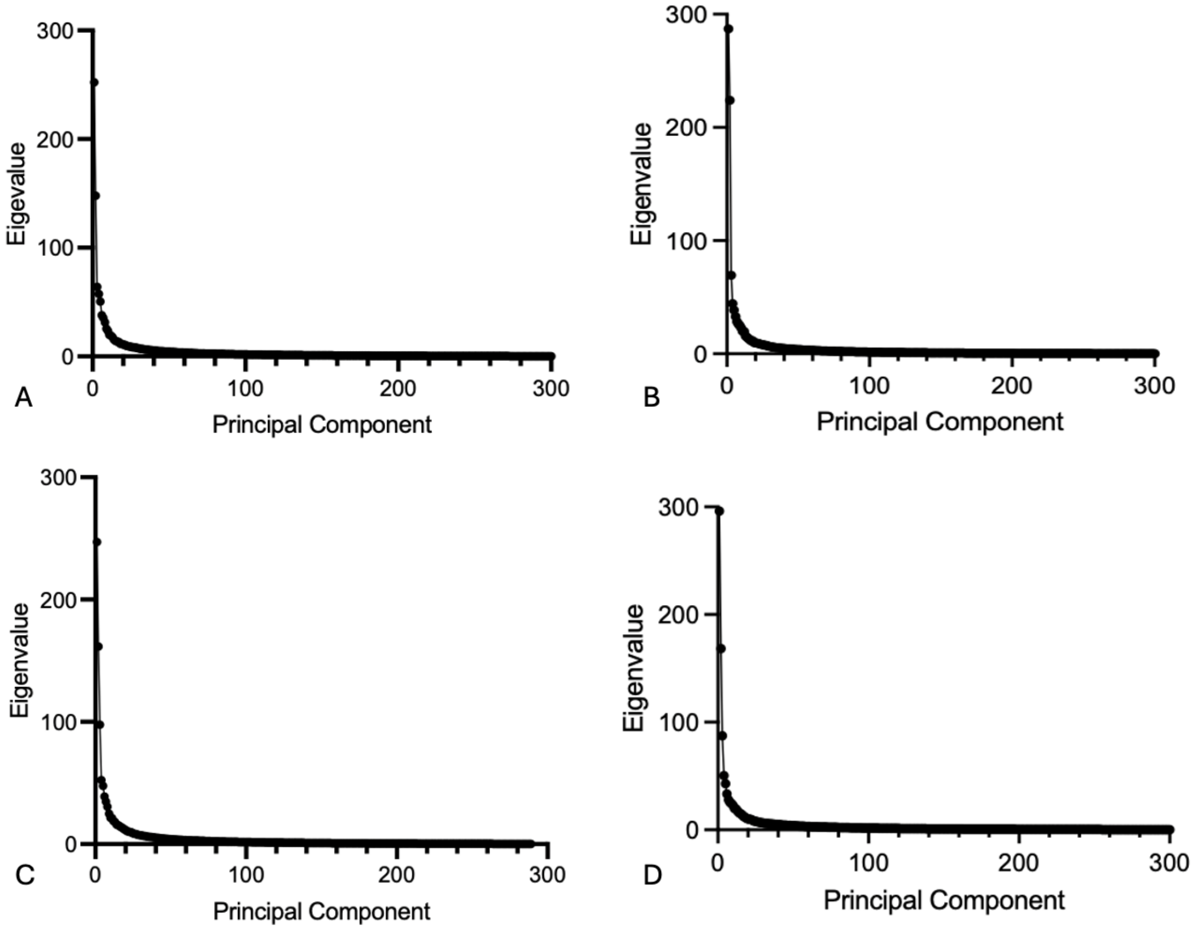


**Figure S1. Scree plots for wild type tyrosinase and mutant variants are shown.** Panel A: WT Tyr. Panels B, C, and D represent the plots for mutant variants R217Q, R402Q, and R217Q/R402Q, respectively.

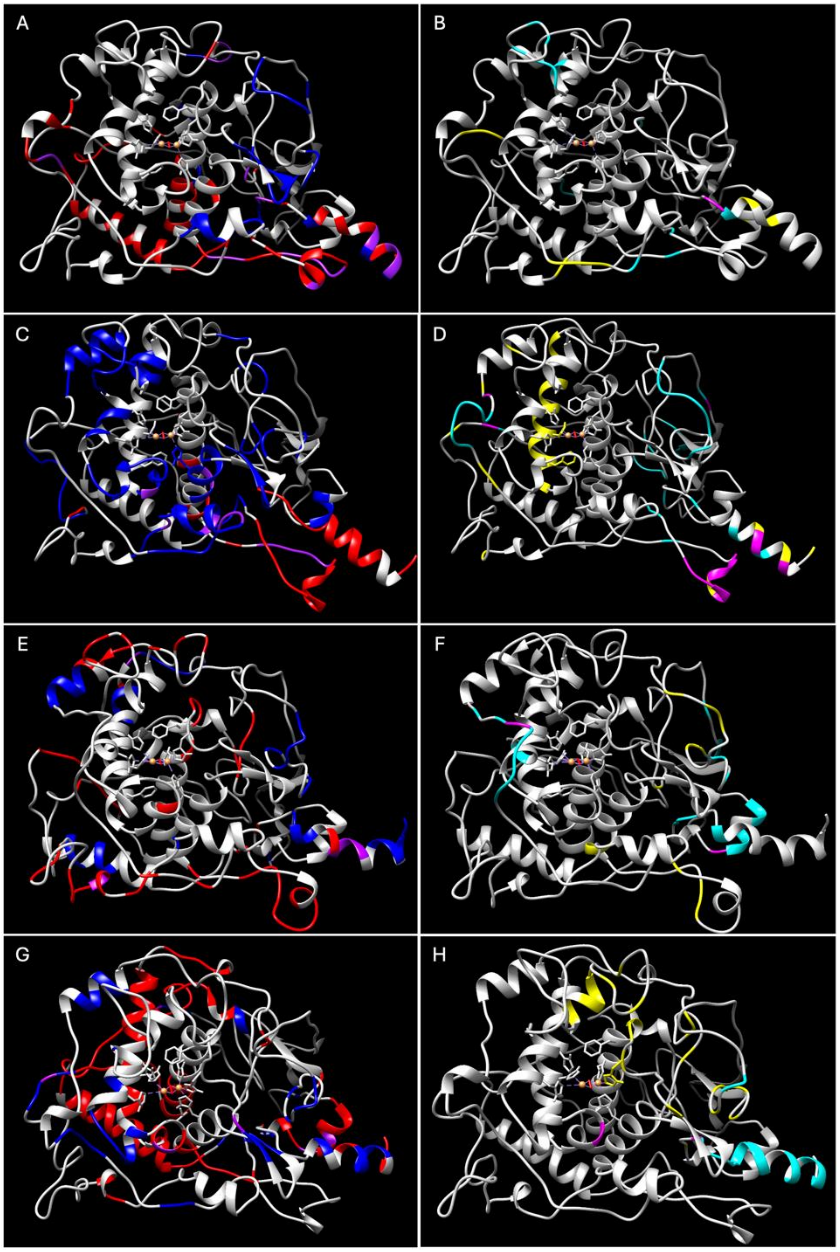


**Figure S2. Representation of PC1 and PC2 subspace for tyrosinase and mutant variant ribbon structures**. Panels A, B: WT tyrosinase, Panels C, D: R217Q, Panels E, F: R402Q, and Panels G, H: R217Q/R402Q. The positive, negative, and overlapping loadings of the PC1 subspace are colored in blue, red, and purple, respectively. The positive, negative, and overlapping loadings of the PC2 subspace are colored in cyan, yellow, and magenta, respectively.

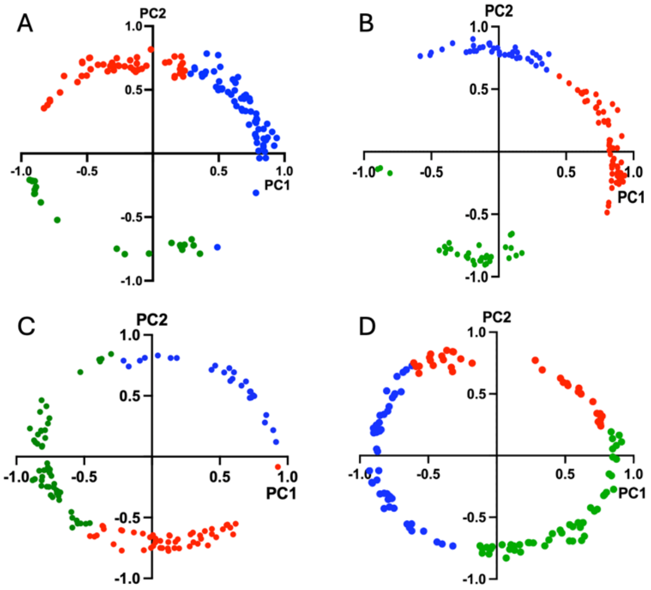


**Figure S3. The graphs obtained for the top 10% of loadings per cluster are shown**. Panel A: WT, Panel B: R217Q, Panel C: R402Q, and Panel D: R217Q/R402Q. This was determined by calculating the distance from the origin and taking the top 10% distance magnitude values.

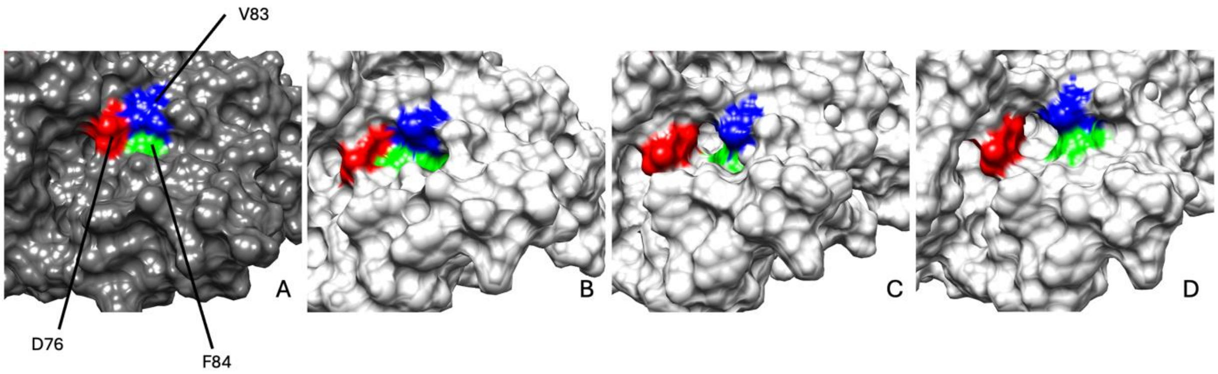


**Figure S4. Instability in R217Q/R402Q mutant variants stems from the disruption of interactions**. Disrupted interactions are between D76 (red), V83 (blue), and F84 (green). The D76-V83-F84 junction is shown for tyrosinase (Panel A, dark gray) and R217Q/R402Q timeframes (light gray) 25 ns (Panel C), 45 ns (Panel D), 85 (Panel E).

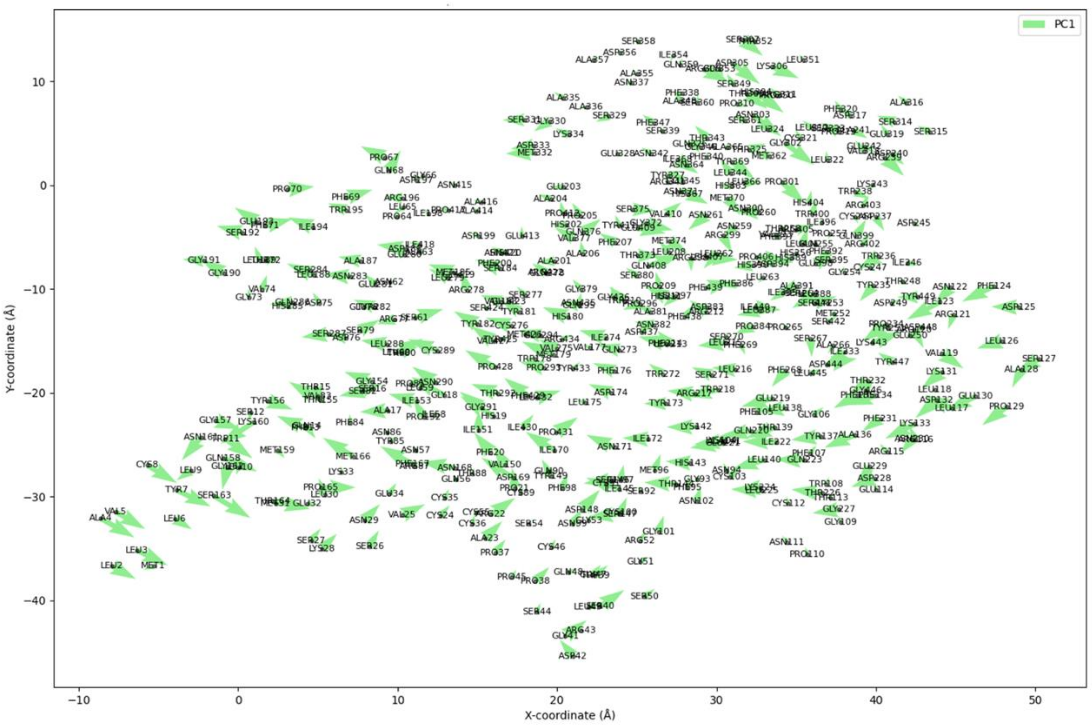


**Figure S5.** PC1 porcupine plot for WT with residue identifiers.

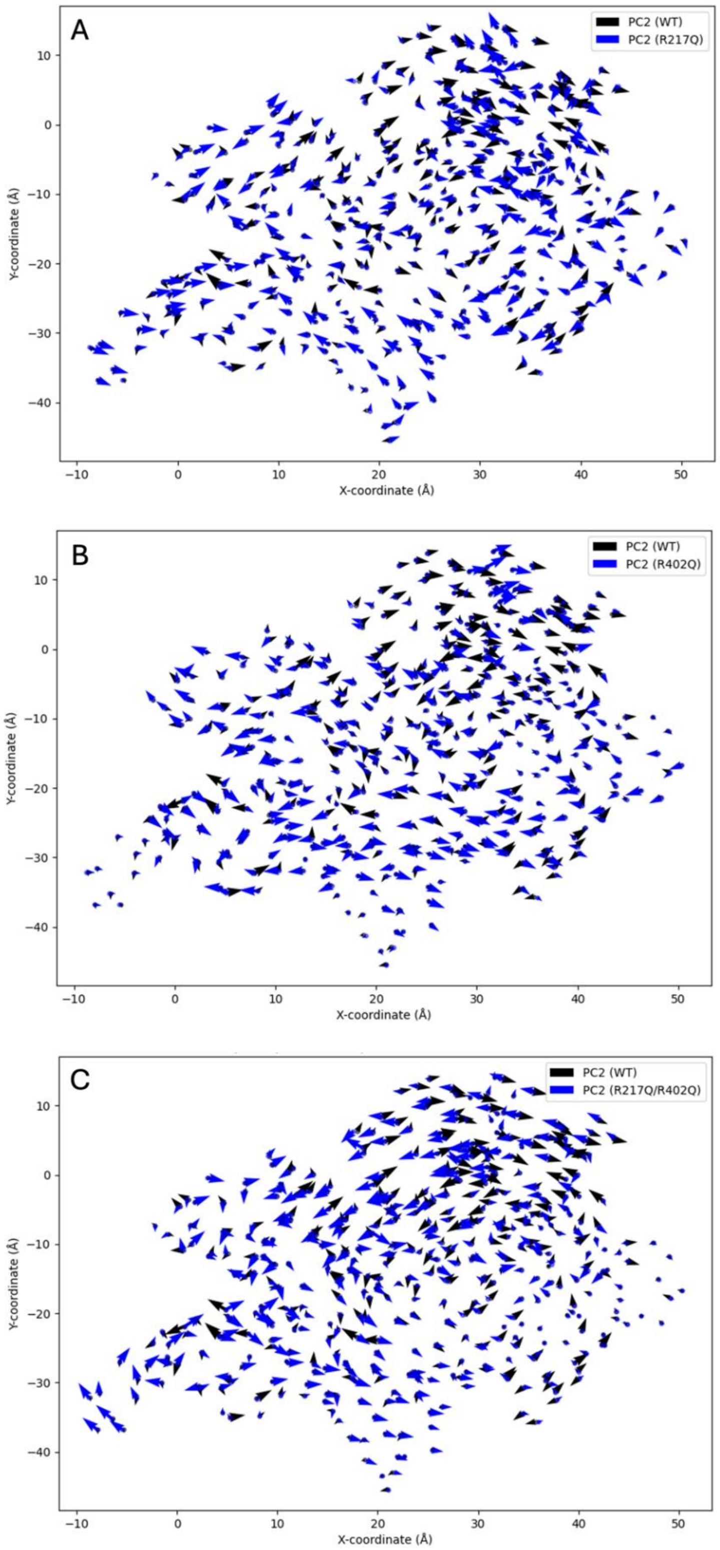


**Figure S6. The superimpositions of PC2 porcupine plots.** Panel A: WT Tyr and R217Q, Panel B: WT Tyr and R402Q, Panel C: WT Tyr and R217Q/R402Q mutant variants.

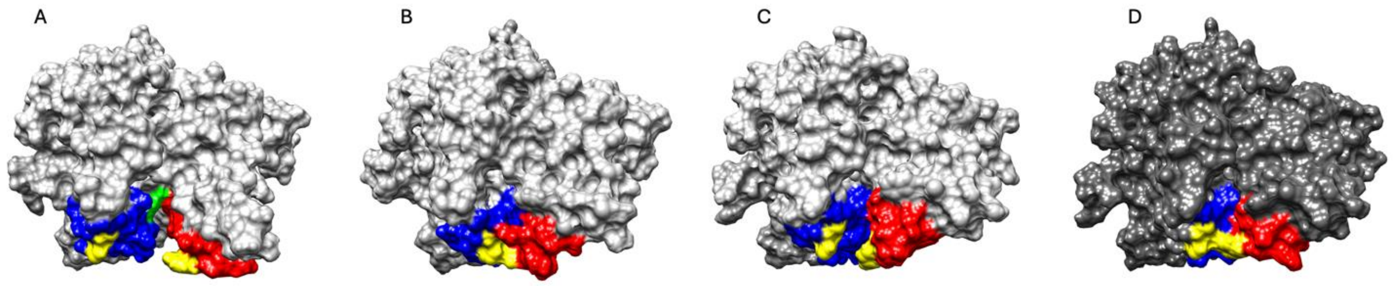


**Figure S7. Instability in R217Q stems results from the loss of interactions**. The interactions are lost between the V150-I170 loop (red), W178 (green), and Q273-E294 loop (blue). The surface renditions proximal to W178 for Clusters 1-3 (Panels A-C) and WT (Panel D) show stability variation amongst the mutant subspaces. All clusters were obtained using the 75 ns timestamp.

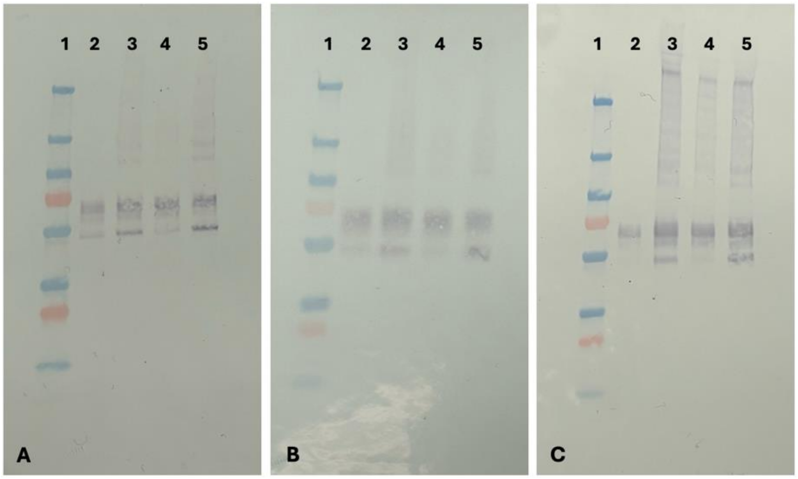


**Figure S8**. **Western blots of tyrosinase and mutant variants**. The blots were obtained using T311 (Panel A), H-109 (Panel B), and C2-85 (Panel C) monoclonal antibodies. For each blot lanes 1-5 are as follows: (1) molecular weight marker, (2) WT tyrosinase, (3) R217Q mutant variant, (4) R402Q mutant variant, and (5) R217Q/R402Q mutant variant. All lysates for each Western blot were run at 0.5 mg/mL concentrations.
